# Comprehensive Analysis of RFID Performance of the iID®BEEscience system

**DOI:** 10.1101/2025.06.03.657638

**Authors:** Jonny Magnay, Ana Paula Cipriano, Sam Macro Versteeg, Francisca Segers, Christoph Grüter

## Abstract

Radio Frequency Identification (RFID) technology has been widely used to study the activity and behaviour of various organisms, including bees. In this study, we rigorously evaluate the performance of a recently developed RFID system (iID®BEEscience from MicroSensys) in tracking bees under different conditions. We assessed the system’s accuracy in detecting and identifying travel direction both in controlled laboratory settings and with free-flying bees. Additionally, we analysed the sources of incorrect or redundant readings and provide a Python script designed to filter out erroneous data, summarising the results in biologically relevant measurements (foraging trip number, trip duration, instances of drifting). Our controlled laboratory tests revealed that the RFID system’s accuracy ranged from 93% to 100% in most cases, though accuracy diminished when a large number of transponders passed through the reader simultaneously. We also found certain transponder positions in relation to the reader position to be less reliable. In field conditions, we tracked 33 foraging trips of RFID-tagged honeybees trained to a sucrose solution feeder and observed a 100% success rate in identifying all foraging trips via the RFID data. Given that the transponders weigh only 2.1 mg, we propose that this system is a reliable tool for studying smaller bees (and other insects), which are currently understudied using RFID technology, although its application to species smaller than honeybees (*Apis mellifera*) requires further testing. This research demonstrates the potential of RFID technology for advancing the study of insect behaviour under field conditions beyond traditional models like honeybees or bumblebees.

## 1. INTRODUCTION

Radio Frequency Identification (RFID) technology is a useful tool in studying animal behaviour, particularly in social insects such as the European honeybee (*Apis mellifera*), where it facilitates the tracking of individual behaviour and colony dynamics. Up until the early 2000s, researchers relied primarily on direct observations and video recordings to study social insect behaviour, identifying individuals using number tags or paint (von Frisch, 1967; Seeley, 1995; Nunes-Silva *et al*., 2019). These methods are labour-intensive and limited by the need for individuals to be visually distinguishable. More recent methods, such as RFID and QR (Quick Response) code technology overcome these issues by allowing the simultaneous recording of multiple individuals, as well as the unlimited number of identification codes and greatly expanded sampling times allowing the collection of a greater volume of data (reviewed in Nunes-Silva *et al*., 2019). The ability of RFID technology to be constantly active also improves detection of rare events, such as honeybee drifting behaviour, as these may be missed by the limited sampling time of direct observations. However, whilst visual obstructions in the form of hive residue and conspecifics may obscure QR tags from view and prevent data recording, RFID radio signals can pass through substances such as glue, wood, and plastic (Decourtye *et al*., 2011), making them a better alternative to study certain questions. However, RFID technology is costly compared to direct observations, detection errors are common, and RFID technology has a limited ability to record other aspects of bee behaviour such as resource transport. Whilst the latter can be resolved using supplementary video recordings of individual honeybee behaviour (Stelzer *et al*., 2010), its high cost may make RFID inaccessible to less well-funded studies.

The number of studies utilizing this technology is extensive, spanning a wide range of species and research questions. It has facilitated studies of rodent social behaviour (Howerton, Garner & Mench, 2012; Sabol, Solomon & Dantzer, 2018), hummingbird foraging behaviour (Bandivadekar *et al*., 2018), and marine crustacean burrow-emergence rhythms (Aguzzi *et al*., 2011), as well as studies seeking to improve livestock management (Brown-Brandl & Eigenberg, 2011; Maselyne *et al*., 2016). In social insects, developments in RFID technology have been especially valuable for research on honeybee foraging (Schneider *et al*., 2012; Feltham, Park & Goulson, 2014; Colin *et al*., 2019; Takahashi *et al*., 2019), homing (Pahl *et al*., 2011), and mating behaviour (Ayup *et al*., 2021; Heidinger *et al*., 2014). It has also helped to further elucidate the mechanisms behind intraspecific reproductive parasitism in large-bodied stingless bees (Van Oystaeyen *et al*., 2013), and their long-distance flight and homing behaviour (Nunes-Silva *et al*., 2020; Costa *et al*., 2021), as well as conopid fly parasitism in bumblebee foragers (Malfi *et al*., 2018). However, whilst there is a large amount of research on corbiculate bees (family *Apinae*), RFID studies targeting other social *Hymenoptera,* such as wasps and ants, remain sparse, with only a few studies investigating decision-making (Robinson *et al*., 2009a; Robinson *et al*., 2009b) and task allocation (Robinson, Feinerman & Franks, 2009) in rock ant (*Tempothorax albipennis*) colonies, drifting in the paper wasp *Polistes canadensis* (Sumner *et al*., 2007), and forager longevity in common wasps (Santoro, Hartley & Lester, 2019). This may be partly due to the large relative size and weight of many RFID tags limiting their use in smaller insects by potentially impacting a variety of life-history traits and reducing the reliability of acquired data (Batsleer *et al*., 2020).

Building on this extensive use of RFID technology in social insects, the present study aims to assess the performance and accuracy of an RFID system recently developed by Microsensys (Microsensys, n.d.) designed to detect passive RFID transponders (mic3®Q1.6 model) weighing just 2.1 mg (Figure 1) with the aim of improving honeybee behavioural research, and enabling further research in smaller social insects and central-place foragers under field conditions, including many smaller bees (e.g. many stingless bees and solitary bees), wasps and ants. Notably, this system possesses a reader containing two antennae, allowing detection of direction using a single reader, and eliminating the need for multiple readers as in older systems (see Fig. 2 in Tenczar *et al*., 2014). The primary objective of our study was to evaluate the RFID system’s performance under controlled laboratory and field conditions and to identify potential sources of error that may impact data integrity. By examining factors that could affect system accuracy, we tried to understand the system’s optimal operating conditions and limitations. This involved testing for reductions in system accuracy caused by each variable, including the Multiple Protocol Command (MPC) cycle (interval in which the system reads and logs the signals received by the antennas) length, transponder speed, orientation, and position relative to the reader, and the effect of varying the distance between two readers, as the manufacturers guide specifies maintaining a minimum of 150cm between two readers. Other factors including the number of transponders simultaneously pulled through the reader, the individual reader used, the effect of water, and the addition of visual tags and acrylic paint have also been tested. A preliminary aim of our study was to investigate the origin of ‘additional readings’ in the data output file, which potentially reduced data usefulness and reliability. To explain this phenomenon, we propose 3 types of errors—Type A, B, and C—and have formulated hypotheses to test these ideas.

**Figure 1:**
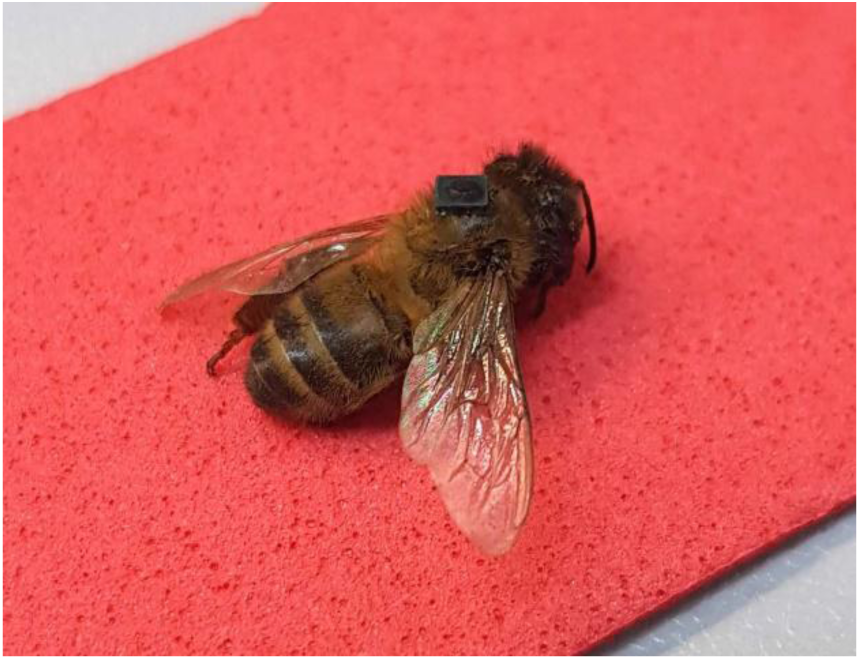
Dead honeybee with 2.1mg mic3®Q1.6 RFID transponder.

**Figure 2:**
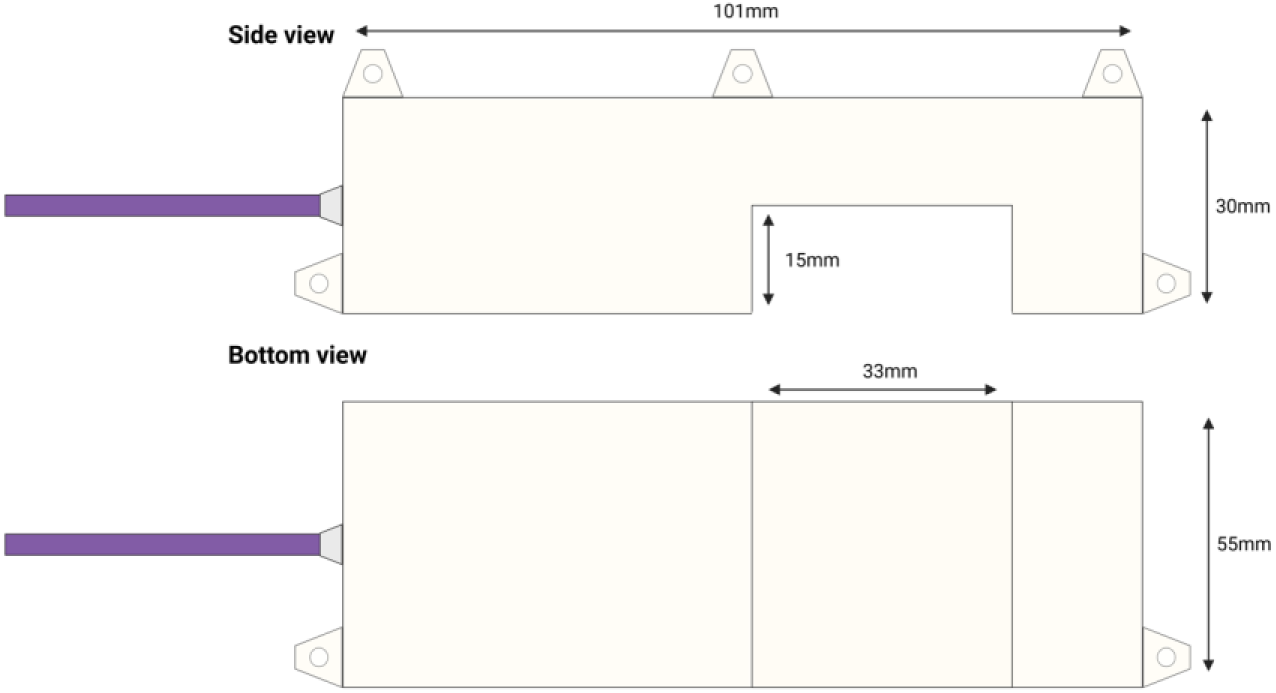
Microsensys iID®science Reader AEB-03.C2D. The Reader casing is made of plastic; holes to attach the Reader to the hive entrance are provided.

These analyses will evaluate the suitability of the Microsensys mic3®Q1.6 RFID system for studying bee behaviour and will facilitate further investigation into the behaviour of honeybees and other central-place foragers. We will discuss the causes behind ‘additional readings’ and evaluate the system’s functionality under laboratory conditions. Additionally, a field test of the system is conducted. Finally, we provide a Python script designed to clean and pre-process the data generated by the system (https://github.com/BristolBeeGroup/BeeDrifting).

## 2. METHODOLOGY

### (a) Apparatus

Our study analysed the performance of a Radio Frequency Identification (RFID) system manufactured by Microsensys. The transponders used were passive RFID transponders of the model Mini-Transponder mic3®Q1.6, measuring 1.6 x 1.6 x 0.4 mm³ with an average mass of 2.1mg. These transponders operate through activation by an electromagnetic field generated by the fixed RFID reader, which powers the transponder, enabling it to transmit data back to the reader for recording. The model of reader used was the iID®science reader device AEB-03.C2D, measuring 101 x 55 x 30mm (Figure 2). Data management and storage were handled by the IID®controller, model CCO-01DC, with 1GB of RAM and 2 antenna outputs, facilitating directional measurements. Data were stored using the BeeID storage software on 16GB USB drives, and power was supplied to the controller from the mains through a 12V power cord, and a PCAN cable and PCAN T-connector connected the iID®BEE controller to the reading devices. In addition to the RFID system, standard number tags (used for queen marking) with an average mass of 1.66mg were used, as well as dead bees obtained ethically from controlled test hives during scheduled observation. In later trials, painting the RFID tags resulted in an average mass of 2.28mg.

### (b) Experimental setup

We conducted the laboratory components of this study over 14 days under constant laboratory conditions, with the iID®science readers positioned 160cm apart on a laboratory bench (Figure 3), exceeding the minimum distance of 150cm suggested by the manufacturers. The IID®controller was positioned equidistant between the readers, which were oriented in an upright position unless specific tests required variations in transponder placement. Field tests with free-flying bees took place between the 13th and 15th August 2024, during which conditions averaged 20°C and sunny.

**Figure 3:**
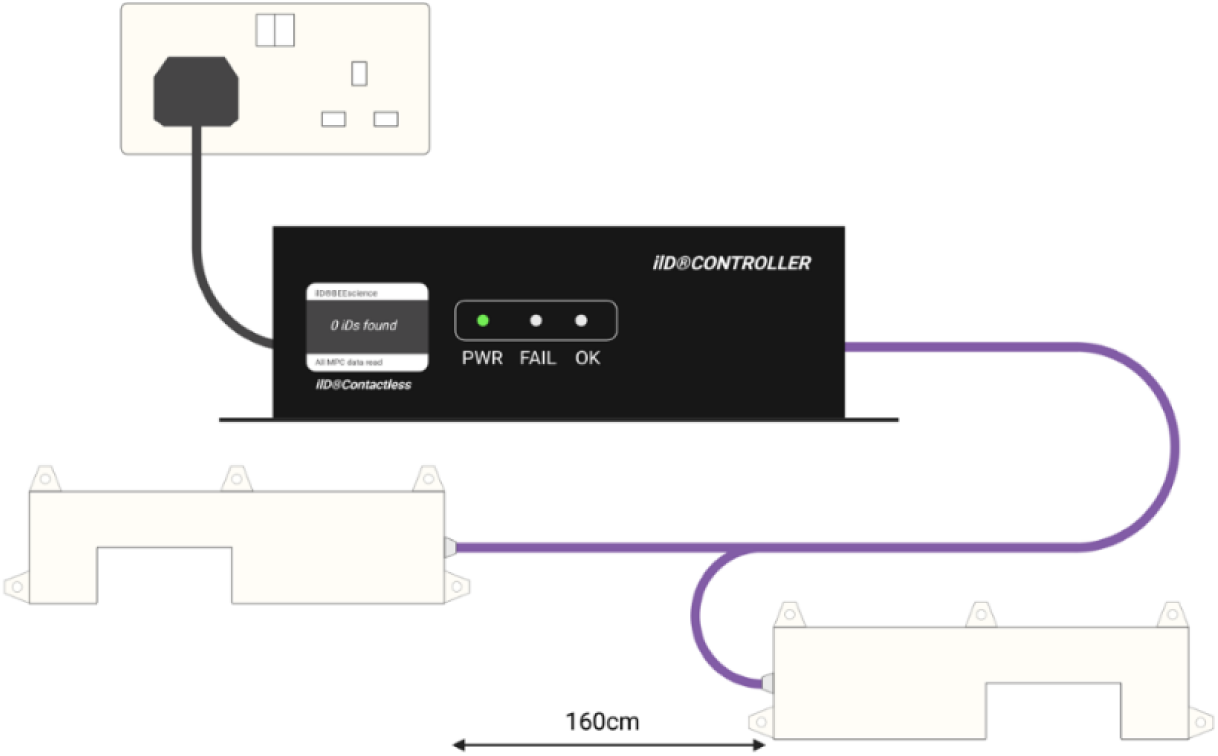
Experimental setup of the RFID system: The setup includes the iID®controller positioned centrally between 2 iID®science readers 160 cm apart. The controller is connected to the readers via PCAN cables. The mains power supply for the controller is also shown.

Data collection settings were standardized to maintain consistency across experiments. Each experiment utilized a 15-second MPC (Multiple Protocol Command) cycle to control for potential variation in accuracy and to minimise the number of additional readings obtained (see section c). Data were recorded in comma-separated file format (CSV), facilitating ease of analysis and compatibility with statistical software packages. Additionally, a daily schedule of controller activity between 04:00 and 00:00 was set up to minimize disruption due to system resetting. For a detailed list of settings, see Figure 4.

**Figure 4:**
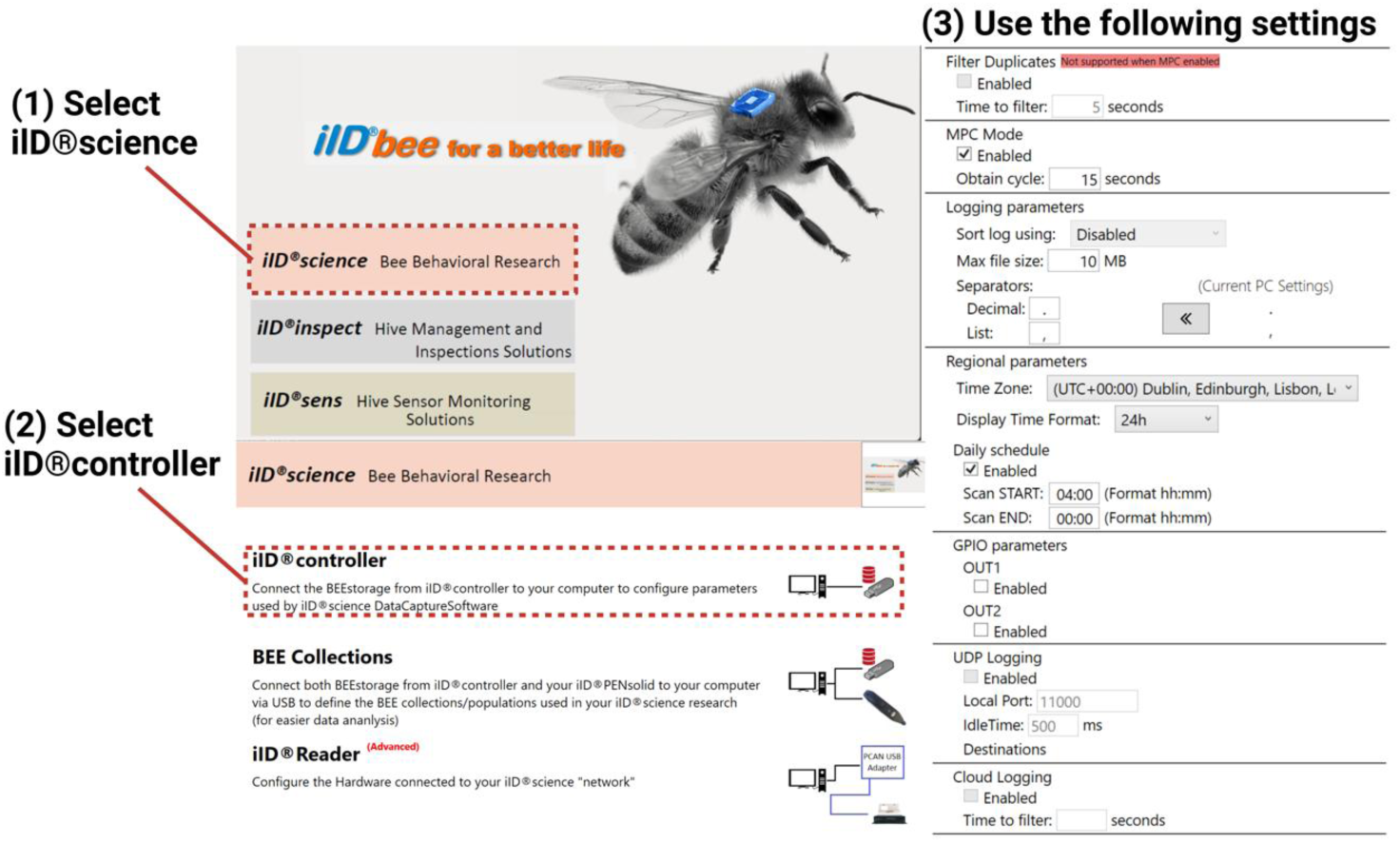
RFID system setup guide.

### (c) Analysis of additional reading errors

A key feature of an accurate RFID system is its ability to classify the direction of a transponder’s movement (e.g., an arriving or departing bee). In preliminary tests, when we simulated ’foraging trips’ by moving transponders in both directions, we observed that the IID® controller not only identified “arriving” and “departing” movements but also classified some as “unknown” in the CSV file. In addition, we found instances where 2 arrivals or 2 departures were reported in very short sequence. Understanding these instances is important to make full use of the collected data.

The IID®controller records data in a cyclical manner defined by the MPC cycle length, specified in the BeeID software, grouping all data collected within this period into a single point recorded in the csv file. This cyclic nature can result in additional readings being recorded exactly one cycle length after (type A error) or before (type B error) an accurate transponder reading. These additional readings pose an issue to the usefulness of the data, and understanding their origin is essential for correctly addressing these errors.

Based on preliminary observations, the following mechanisms for the origin of additional readings have been proposed. These additional readings appeared to occur during the transition between one cycle ending and another beginning while the transponder was still within range of one or both reader antennae. In such cases, additional readings often appeared as ’unknown’ in the csv file when recorded by only one antenna or appear directional (2 “arriving” or “departing” in short sequence) if recorded by both. To give an accurate reading with no additional readings, the transponder must fully pass through and out of the ranges of both antennae during one cycle duration (Figure 5a shows an example of an “arriving” transponder movement). Type A errors occurred when the cycles turnover whilst the transponder is still in range of the second antenna, generating an ‘unknown’ reading exactly one cycle duration after the directional reading (Figure 5b). Type B errors occurred when the cycles turnover whilst the transponder is only within range of the first antenna and has not yet entered the range of the second antenna, giving an ‘unknown’ reading exactly one cycle duration before the directional reading (Figure 5c). We also observed occasions when the cycles turn over whilst the transponder is in range of both antennae, giving a second directional reading and generating a type C error (Figure 5d). In this case, we suggest the first value should be considered correct. Whilst Figure 5 shows only ‘arriving’ errors, these errors also occur for ‘departing’ transponders (Figure 6).

**Figure 5:**
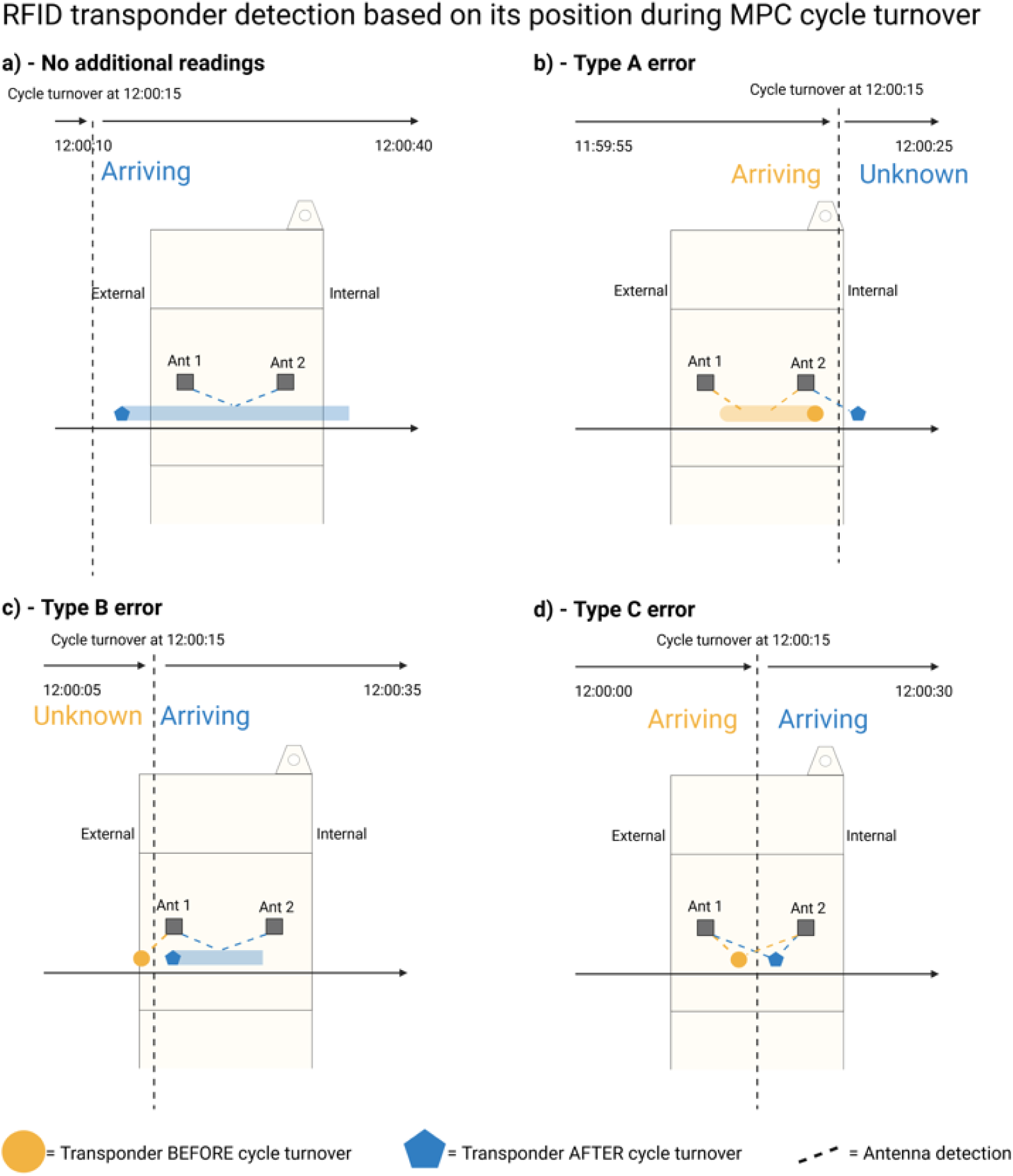
Possible outcomes of RFID ‘arriving’ transponder detection depending on its position relative to each antenna at the point of a 30-second cycle turnover at 12:00:15: The same errors also occur for ‘departing’ insects. Detected movements before cycle turnover are shown in large orange text, with those detected after shown in blue. The position of the RFID transponder before cycle turnover is represented by an orange circle, and by a blue pentagon after cycle turnover. The path of the transponder is depicted using blue and orange lines. (A) After one cycle turnover and before a subsequent turnover, the transponder passes through and out of the range of both antennae, giving a single directional reading. (B) At the point of turnover, the transponder has passed both antennas and remains within range of antenna 2, giving a type A error. (C) At the point of turnover, the transponder is in range of antenna 1 but not yet antenna 2, giving a type B error. (D) At the point of turnover, the transponder is in range of both antennas, giving a type C error.

**Figure 6:**
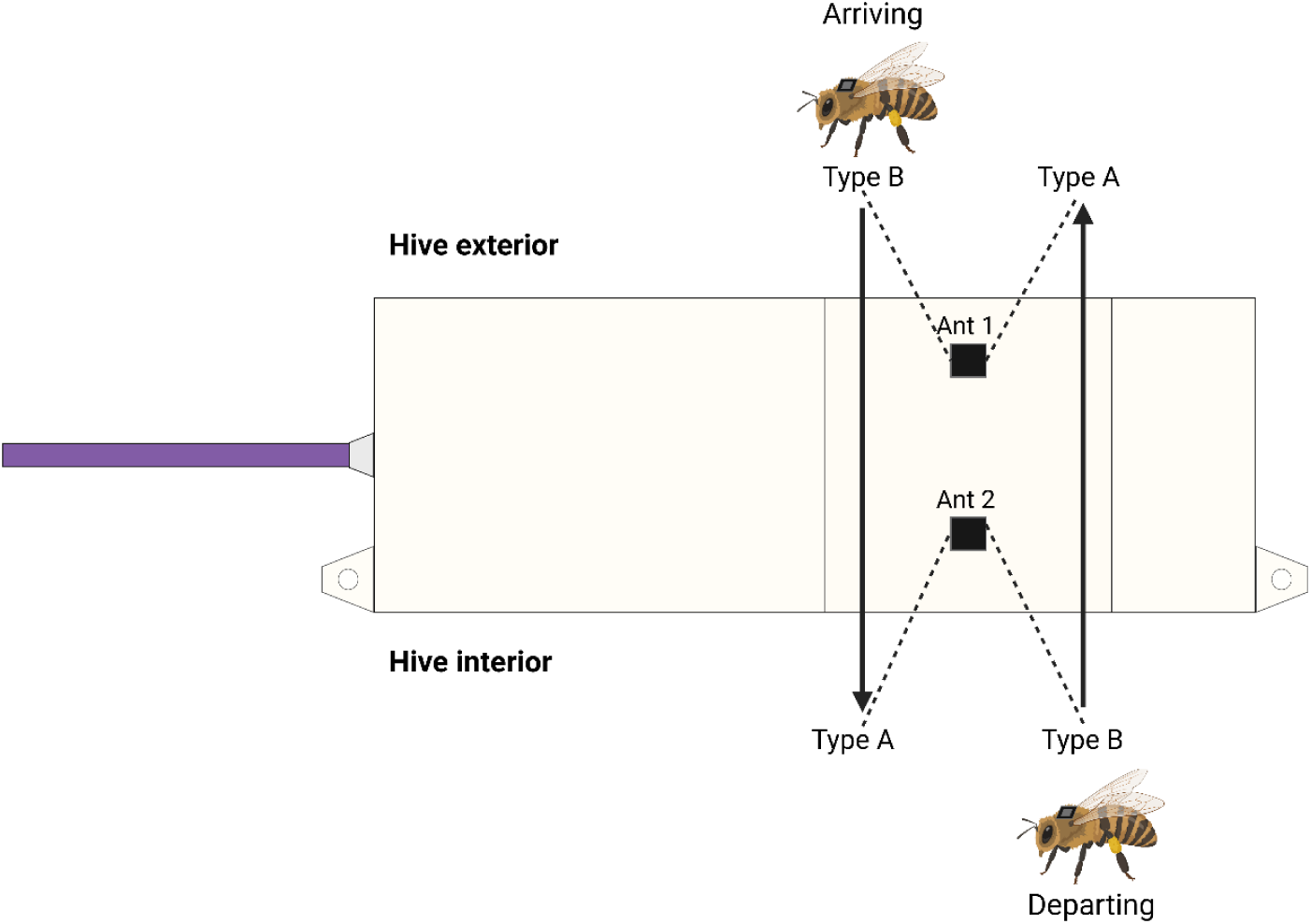
Illustration of how the ’unknown’ additional reading corresponds to specific antennas based on the type of error: (A) Type A Error: An unknown reading corresponds to the antenna that the transponder encountered second. (B) Type B Error: An unknown reading corresponds to the antenna that the transponder encountered first. The diagram depicts the sequence of encounters and the logical association of the ‘unknown’ reading with the respective antenna.

To investigate whether these ideas are likely correct, the following predictions have been set out:

1. When the length of the MPC cycle is reduced, the overall number of additional readings will increase proportionally due to an increased likelihood of the cycles transitioning whilst the transponder is in range of the reader.
2. There will be an equal number of type A and type B errors since the cycle is equally likely to transition on either side of the reader.
3. For type A errors, the ‘unknown’ additional reading will correspond to the antenna encountered secondarily during movement through the reader. For type B errors, it will correspond to the antenna encountered first (Figure 6).

To test these hypotheses, we analysed the type (A, B and C) and frequency of additional readings under varying MPC cycle lengths (5 sec, 10 sec, 15 sec, 30 sec). We placed a single RFID transponder upright on a 3mm-high ‘foam bee’ strip constructed from stacked 1.5mm craft foam sheets (Figure 7), secured it with nail varnish, and pulled it through the reader by hand at ∼30mm/s by timing 1.8 seconds on a stopwatch. This setup mimics the average walking speed (Nouvian and Galizia, 2019) and height (measured from dead workers) of honeybees, ensuring realistic testing conditions and controlling for distance from the sensor and speed of drag. The actual time of movement was noted, allowing comparison with the recorded RFID data. Observer bias was controlled for by covering the controller display so the investigator could not see the detection outcome. The direction of pulls was alternated to facilitate efficient data analysis, with a 65-second gap between each pull to ensure data points were categorized into distinct cycles.

**Figure 7:**
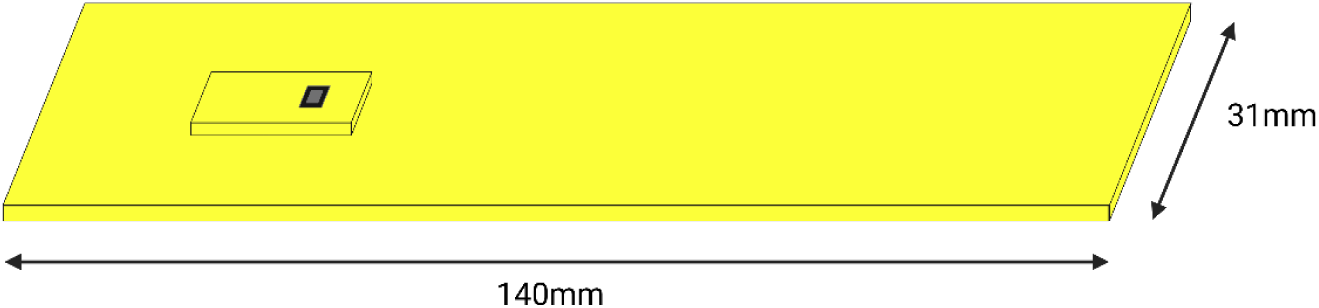
31 x 140mm ‘Foam bee’ strip with single RFID transponder.

Data were analysed using the software R (v4.2.1). For prediction 1, the 15-second MPC cycle was considered a reference group and used to calculate the expected proportions of additional readings for the 5, 10, and 30-second cycles (25%, 12.5%, and 4.167% respectively). A chi-squared test was used to test for significant differences between actual and expected proportions of additional readings for each MPC cycle. For prediction 2, a chi-squared test was used to test for differences in the proportion of type A and type B errors within each MPC cycle. For prediction 3, the antenna recording each type A and type B error was identified and compared to the expected antenna based on our hypothesis (Figure 6). The proportion of matching data points was then calculated.

### (d) Testing system accuracy

To assess the accuracy of the RFID system under various conditions, we designed a series of tests to identify and quantify potential sources of error and inform efforts to improve its accuracy. The dependent variable in all tests was the accuracy rate of the RFID system, defined as the percentage of movements of the RFID transponder through the reader whose time and direction were correctly recorded without erroneous additional readings. Erroneous additional readings are type A, B, or C errors as described in section c, but which show an incorrect direction. For example, for a departure from the hive, a type A, B, or C error may show either ‘unknown’ or ‘departing’, whereas an erroneous additional reading would read ‘arriving’. Additionally, if a transponder movement was not recorded at all, or a movement was recorded when there had not been one, these were also classed as errors. Type A, B, and C ‘errors’ were excluded from the analysis of accuracy as they are independent of system accuracy, instead relating to the cyclic nature of the system (see section 2c).

As with the tests for additional readings, we placed a single RFID transponder upright on a 3mm-high ‘foam bee’ strip constructed from stacked 1.5mm craft foam sheets (Figure 7), secured it with nail varnish, and pulled it through the reader by hand at ∼30mm/s by timing 1.8 seconds on a stopwatch. The actual time of movement was noted, allowing comparison with the recorded RFID data and identification of inaccurate data points. In addition to the ‘foam bee’ strip used for the majority of tests, some tests necessitated the use of a ‘dead bee’ strip to control for any impact of honeybee physiochemistry. This involved a dead honeybee being secured to a foam strip of 1.5mm height and 31mm width using nail varnish (Figure 8). Whilst these strips were suitable for Tests 1-4 and 6-12, **Test 5** required the creation of 13mm-wide strips to assess the effects of side position in the reader (Figure 8).

**Figure 8:**
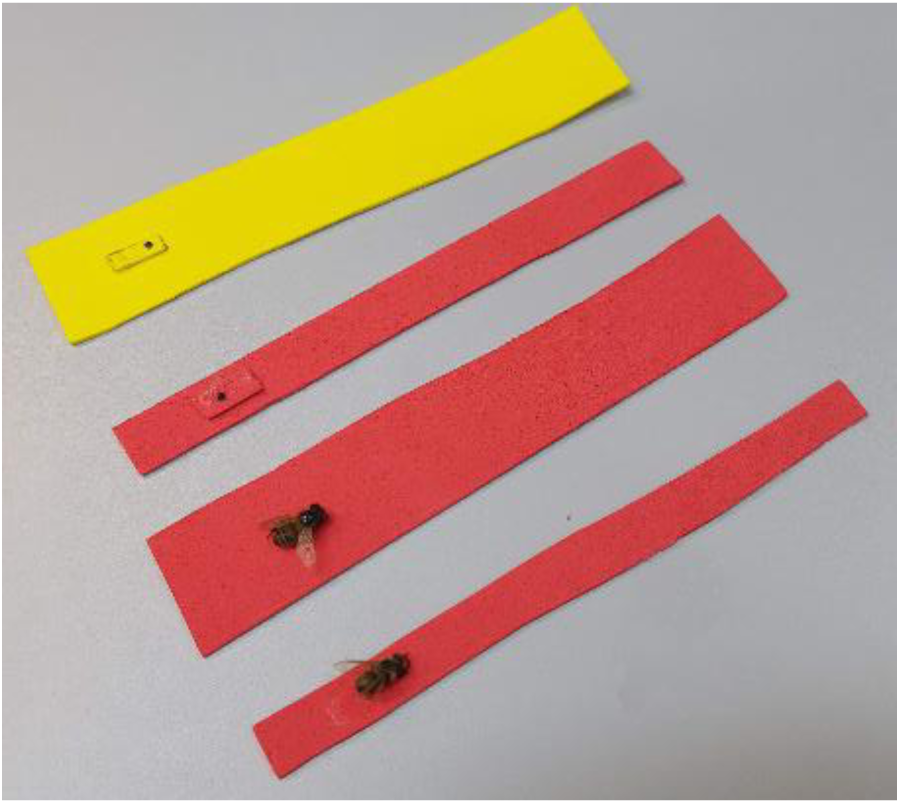
Additional ‘foam bee’ and ‘dead bee’ strips: Top to bottom: 31 x 140mm ‘foam bee’ strip; 13 x 140mm ‘foam bee’ strip; 31 x 140mm ‘dead bee’ strip; 13 x 140mm ‘dead bee’ strip.

The following ‘default conditions’ were utilised across all tests to ensure consistency: a 15-second MPC cycle, providing the greatest accuracy whilst minimising the time spent collecting data; the speed of transponder movement maintained at approximately 30mm/s by pulling it through the reader by hand, timing 1.8 seconds on a stopwatch; 60 reads for each test (30 arrivals and 30 departures); a time interval of 65 seconds between each read to prevent their grouping into the same MPC cycle; consistent transponder placement using the ‘foam bee’ or ‘dead bee’ strips; consistent 12mm distance from the readers antennae; transponders in an upright position; nail varnish used to secure the transponders, mimicking real-world usage; controller display covered to minimise investigator bias. A detailed description of each test follows.

#### Test 1 - MPC cycles

To decide on which MPC cycle to use as a default, we analysed the effects of MPC cycle length (5 sec, 10 sec, 15 sec, 30 sec) on system accuracy. All subsequent tests utilized a 15-second MPC cycle, and the data of the 15-second MPC group from Test 1 is henceforth used as the default setting for which to compare subsequent tests involving the ‘foam bee’ strip.

#### Test 2 – Dead bee

Any impact of a bee’s physical properties on system accuracy was tested by pulling a ‘dead bee’ strip through the primary reader.

#### Test 3 - Speed of movement

We tested the effect of transponder movement speed on system accuracy by pulling the ‘foam bee’ strip through the primary reader at speeds of ∼10mm/s and ∼50mm/s. To reproduce these speeds, a ‘pull-time’ of 5.5 and 1.1 seconds respectively was maintained across the reader. The accuracy of each test was then separately compared to the default speed of ∼30mm/s, the average honeybee walking speed (Nouvian and Galizia, 2019).

#### Test 4 - Transponder orientation

To test the effect of transponder orientation on the bee (upright, upside-down) on system accuracy, we secured a singular RFID transponder upside-down to a ‘foam bee’ strip using nail-varnish (Figure 9A). Identical procedures were applied to a test group of an upside-down transponder on a ‘dead bee’ strip (Figure 9B).

**Figure 9:**
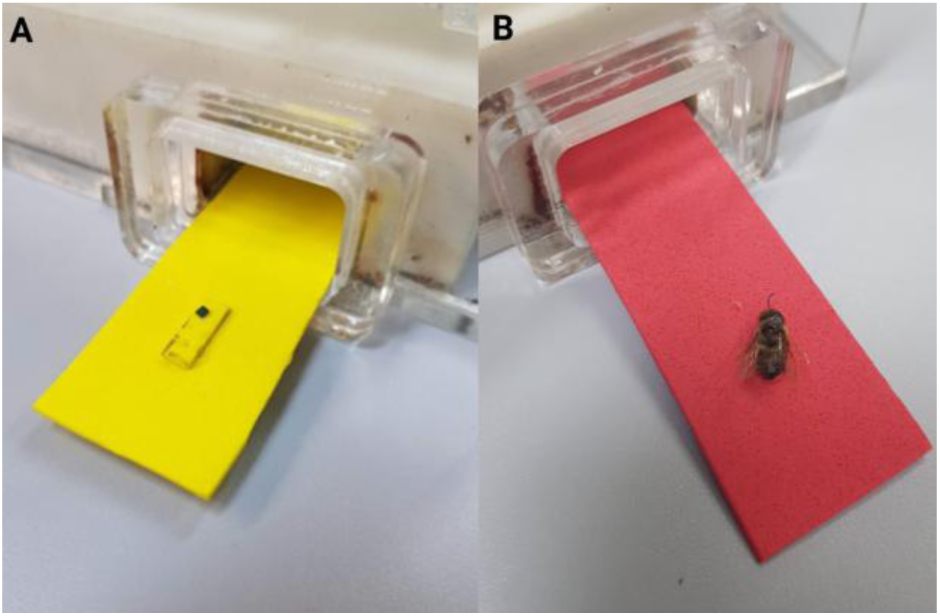
Primary reader setup with upside-down transponders using (A) the 31 x 140mm ‘foam bee’ strip and (B) the 31 x 140mm ‘dead bee’ strip.

#### Test 5 - Position in reader

Testing for any variations in accuracy based on transponder position through the reader (bottom, top, left, right; see Figure 10) was crucial to identify limitations involving transponder positioning, as bees may travel on all entrance surfaces as they enter the hive. To test the top of the primary reader, it was turned upside-down and kept level using strips of foam, and we pulled the ‘foam bee’ strip through at default speed (Figure 11). To test the left side of the primary reader, it was oriented with the cable end facing downwards and was propped up with foam sheets, with a 13mm wide ‘foam bee’ strip pulled through. After highly anomalous results, this side was repeated with the reader positioned so the cord did not touch the lab bench, as it was believed that errors with antenna 2 were caused by the cable bending too far (Figure 11B). To test the right side of the primary reader, it was oriented with the cable end facing upwards (Figure 11C), again pulling the 13mm wide ‘foam bee’ strip through. Identical procedures were followed using ‘dead bee’ strips, which were compared to the reference of the bottom of the reader (Figure 11D, E, F).

**Figure 10:**
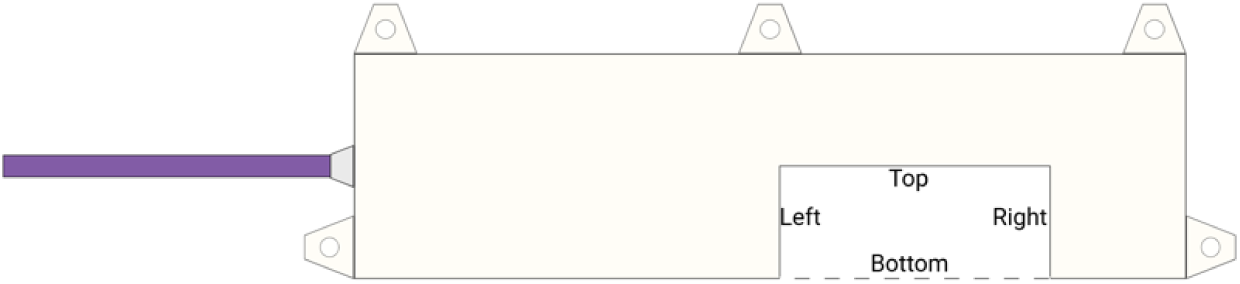
Classification of position in the reader (Top, bottom, left, right)

**Figure 11:**
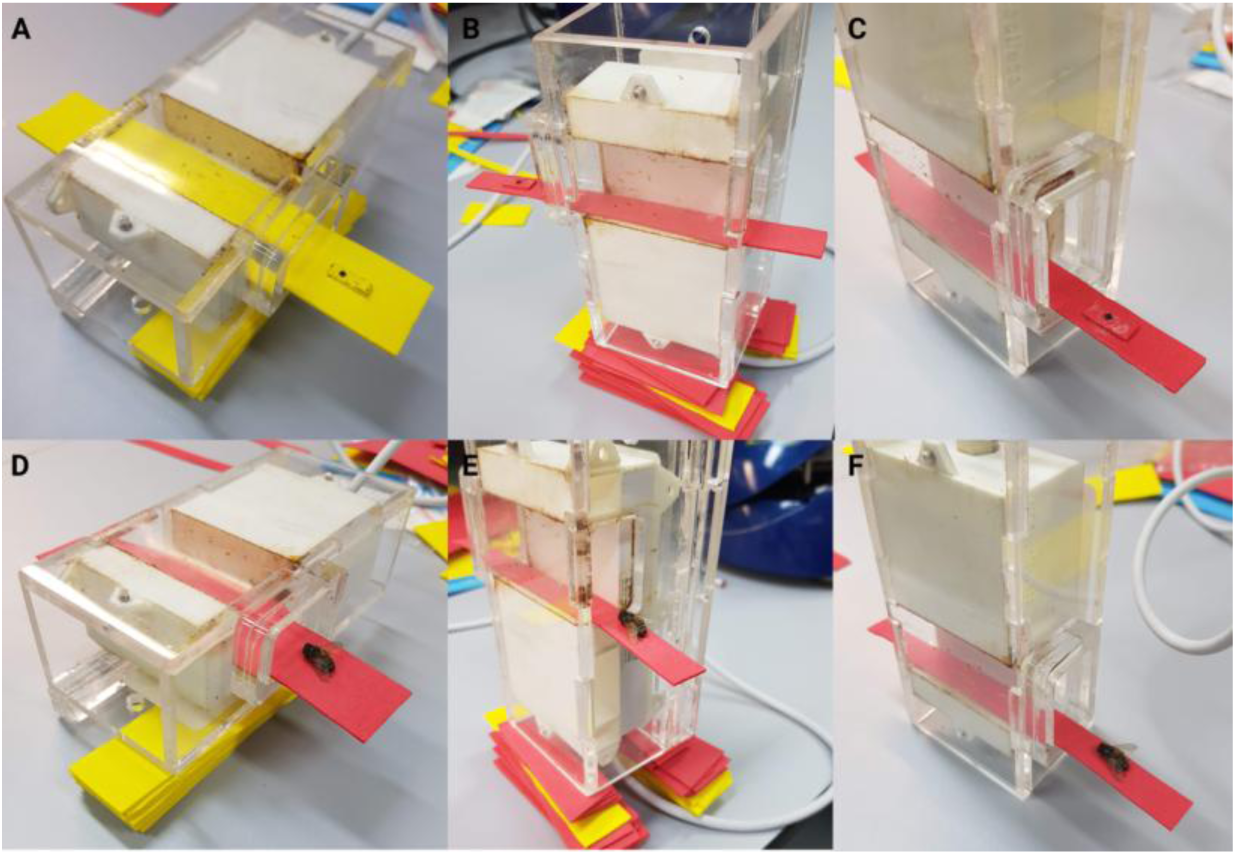
Primary reader setup at different orientations to test position in the reader: (A) Top of the reader using 31 x 140mm ‘foam bee’ strip; (B) Left of the reader using 13 x 140mm ‘foam bee’ strip; (C) Right of the reader using 13 x 140mm ‘foam bee strip; (D) Top of the reader using 31 x 140mm ‘dead bee’ strip; (E) Left of the reader using 13 x 140mm ‘dead bee’ strip; (F) Right of the reader using 13 x 140mm ‘dead bee’ strip.

#### Test 6 - Effectiveness of the second reader

We tested the accuracy of a second reader to ensure it was equal to that of the primary reader. We performed tests with both ‘foam bee’ and ‘dead bee’ strips.

#### Test 7 - Distance from reader

To assess both limitations of the system involving distance from the readers antennae, and the potential for use with hive entrances larger than the readers height, it was necessary to investigate how the distance of a transponder from the reader affected system accuracy. Distance from the second readers antennae was manipulated through stacking 1.5mm thick foam strips to change either the height of the transponder inside the reader (for 9mm distance) or the height of the reader around the transponder (for 15mm and 18mm distance) (Figure 12).

**Figure 12:**
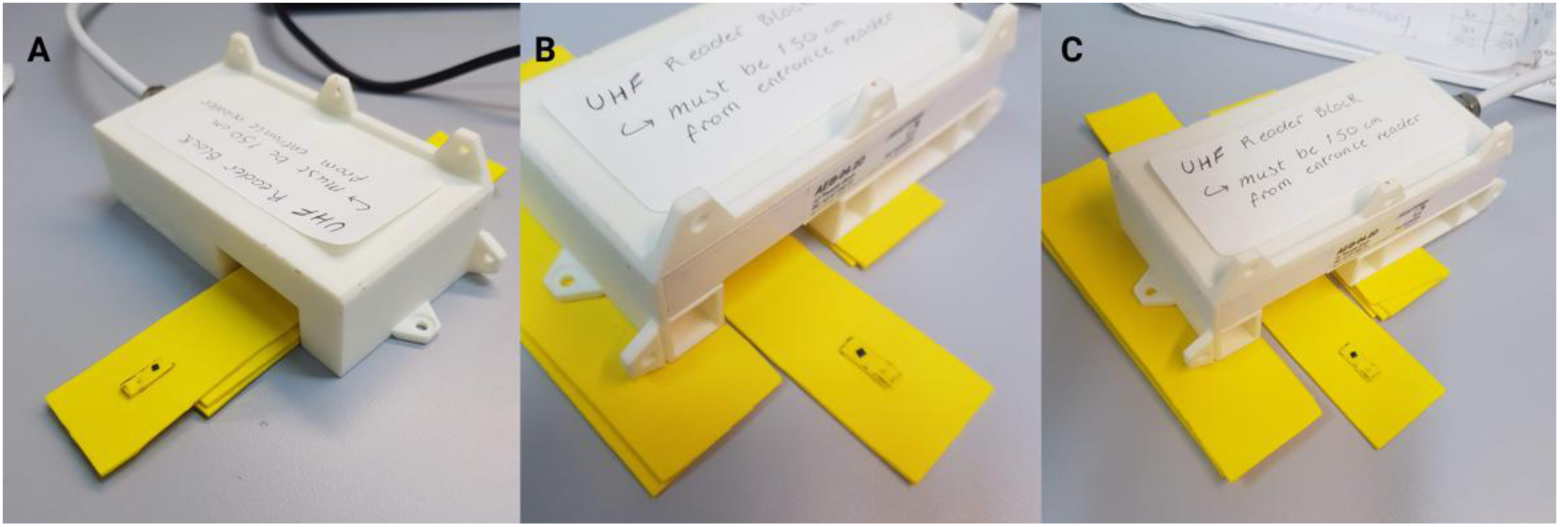
Second reader setup using 31 x 140mm ‘foam bee’ strips: (A) 9mm distance tested using 3mm high ‘foam bee’ strip stacked on 2x 1.5mm high foam strips; (B) 15mm distance tested using 3mm high ‘foam bee’ strip and reader raised on 2x 1.5mm high foam strips; (C) 18mm distance tested using 3mm high ‘foam bee’ strip and reader raised on 4x 1.5mm high foam strips.

We used the ‘foam bee’ strip to assess differences in accuracy of these test groups and the reference group of 12mm.

#### Test 8 - Multiple transponders

During real-world data collection, it is likely that multiple bees possessing RFID transponders enter or depart the hive through the reader at the same time, causing potential signal disruptions. To test whether multiple transponders passing through the reader at once reduces system accuracy, we pulled two test groups, involving 5 or 10 transponders on a foam strip, through the reader (Figure 13). In each treatment the same distance between each transponder was maintained to control for transponder density and ensure that only the number of transponders passing through the reader at once changed.

**Figure 13:**
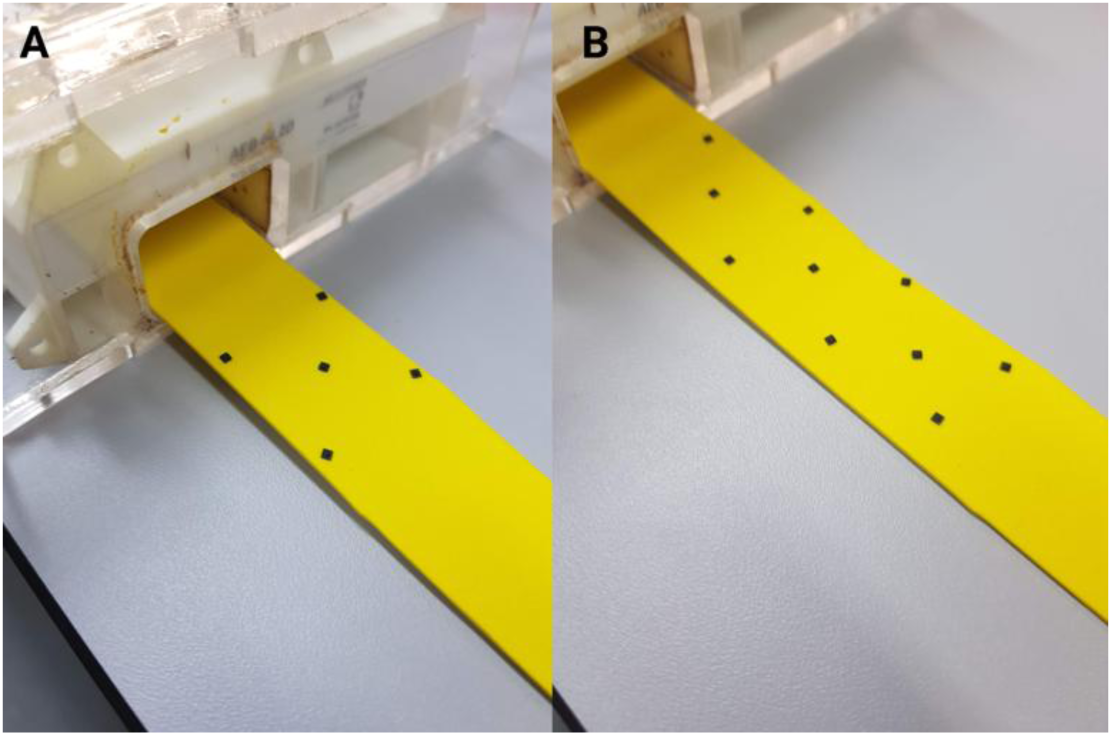
Primary reader setup using the 31 x 140mm ‘foam bee’ strip with (A) 5 transponders, and (B) 10 transponders.

#### Test 9 - Visual tag

An important development in the field of RFID insect tagging would be the simultaneous use of both an RFID transponder and a number tag. The number tag could potentially block the RFID signal, hence we tested whether the addition of a number tag reduces the accuracy of the system. We secured number tags onto the top of the RFID transponder using nail polish, testing accuracy using both ‘foam bee’ and ‘dead bee’ strips (Figure 14).

**Figure 14:**
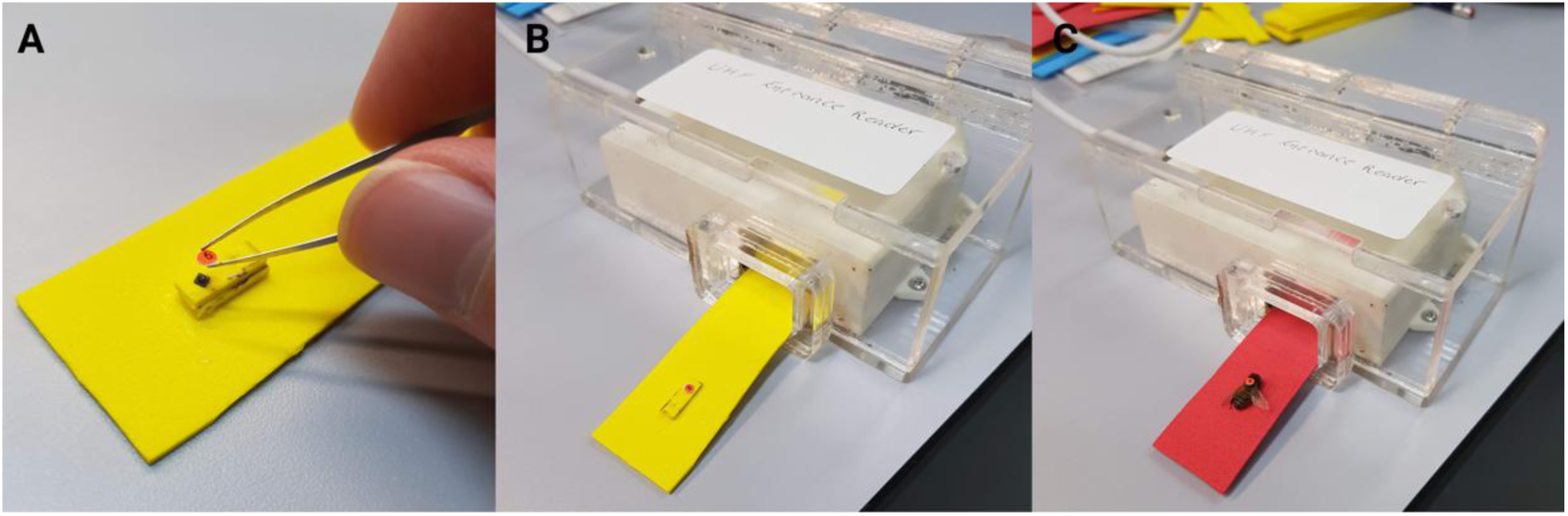
Application of number tag (A), and primary reader setup using the (B) 31 x 140mm ‘foam bee’ strip with an additional visual tag, and (C) ’31 x 140mm ‘dead bee’ strip with an additional visual tag.

#### Test 10 – Distance between readers

To test whether the distance between the readers has an impact on the systems accuracy, we positioned the readers 15cm from each other and pulled a ‘foam bee’ strip through the reader at default speed (Figure 15). Differences in accuracy between this test group and the default distance of 150cm were tested.

**Figure 15:**
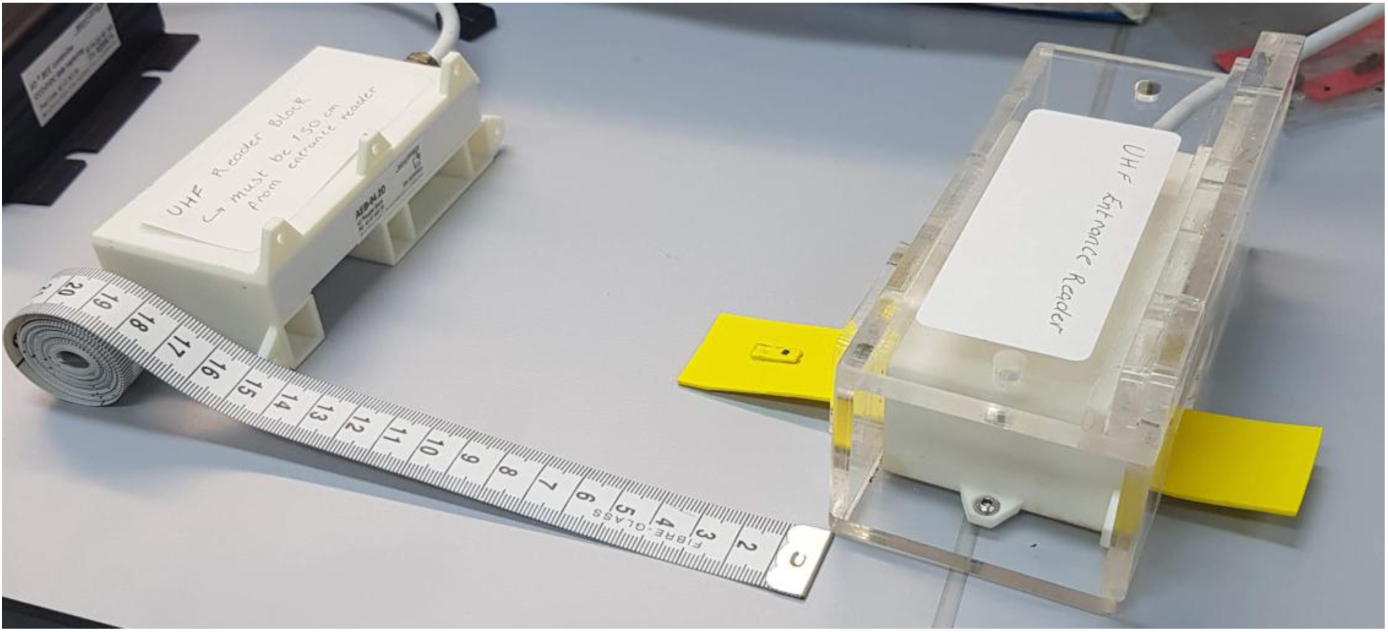
Primary reader setup positioned 15cm from the second reader, using the 31 x 140mm ‘foam bee’ strip.

#### Test 11 - Effects of water

To test the resistance of the transponders to water, we soaked a single transponder in water for 5 minutes, then secured it to a ‘foam bee’ strip and pulled it through the reader. We then compared the accuracy of this test to the corresponding test without water treatment.

#### Test 12 – Effects of painting the transponder

An alternative method of individual honeybee identification involves the use of distinct colour combinations. For this to function experimentally, painting the transponders must not reduce their detection rate. To test this, we painted 5 transponders with acrylic paint (‘docrafts ARTISTE’) (Figure 16), secured them individually to a ‘foam bee’ strip, and pulled them each through the primary reader 10 times.

**Figure 16:**
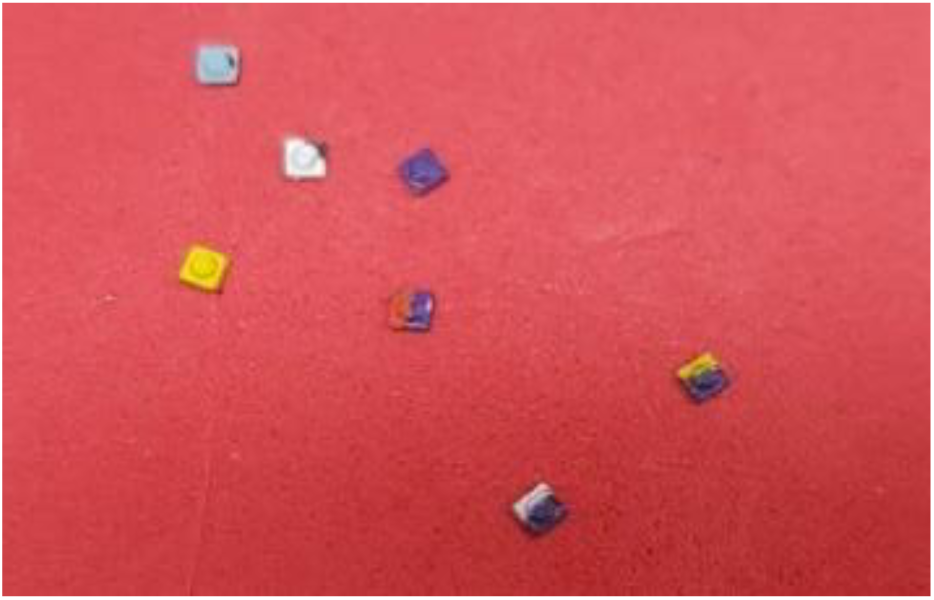
RFID transponder painted with acrylic paint.

### (e) Field testing

To assess RFID system performance in field conditions and collect a sample of honeybee foraging data, the readers were installed onto the entrance of a single observation hive kept in an observation shed using screws and electrical tape (Figure 17A). The hive contained a queen-right colony of *Apis mellifera* with ∼3,000-4,000 worker bees. The IID®controller was positioned on a raised platform between the 2 readers and was plugged into the mains using the 12V power cord and extension lead (Figure 17B). We trained honeybees from a similar, second observation hive to use a sugar-water feeder 150m from the hive using standard training methods (von Frisch 1967) (Figure 17C).

**Figure 17:**
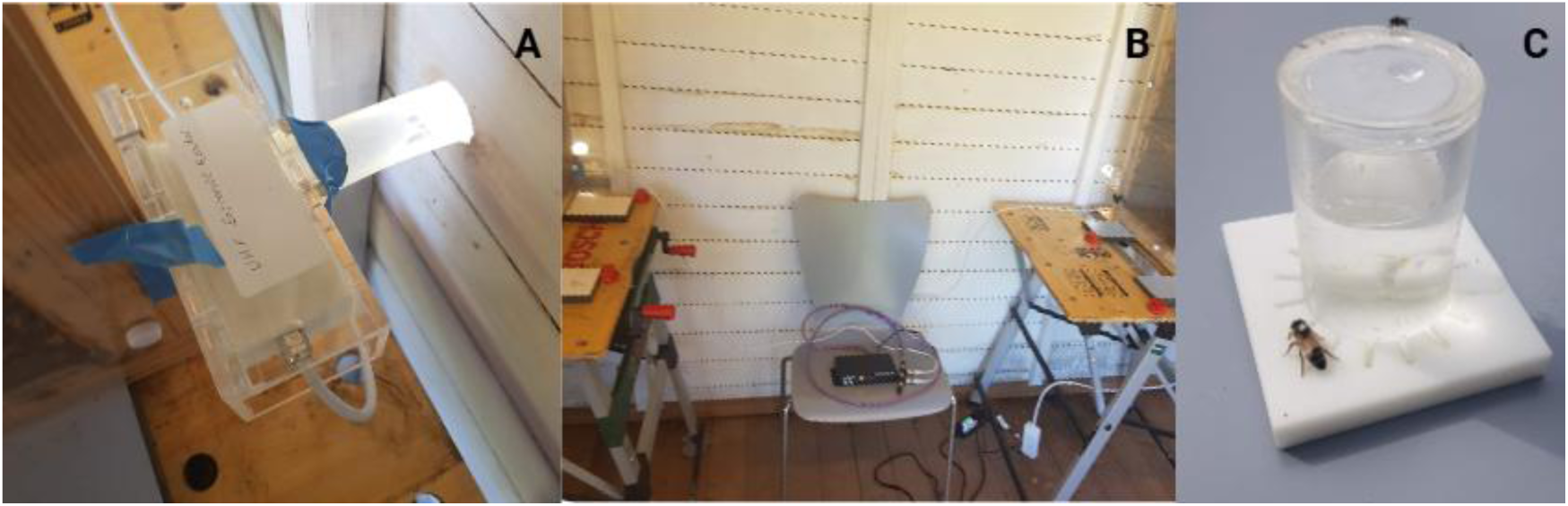
(A) RFID reader installed on observation hive entrance; (B) Observation hives 150cm apart; (C) Sugar water feeder 150m from the hive.

On the 9^th^ August, 9 forager bees found on the feeder were tagged with an RFID transponder and a visual number tag was glued on top of the RFID transponder. However, after 2 days all number-tags had been removed, and individuals could no longer be distinguished. In a second attempt on the 13^th^ of August, 9 forager bees trained to the feeder were tagged with an RFID transponder painted with distinct colour combinations. On the 14^th^ of August, the feeder was set up at 12.30pm and observed between 12:35pm and 14:35pm whilst the RFID controller was active, recording both the time of arrival and departure to the food source through direct observation, and time of arrival and departure from the colony using RFID data. These were then plotted in a line graph of location against time, and direct focal observations were compared with RFID data to assess system accuracy.

## 3. RESULTS AND DISCUSSION

### (a) Additional readings

Overall, 5 second MPC cycles generated the highest proportion of additional readings (20%), followed by 10 (13.33%), 15 (8.33%), and 30 seconds (5%) (Figure 18).

**Figure 18:**
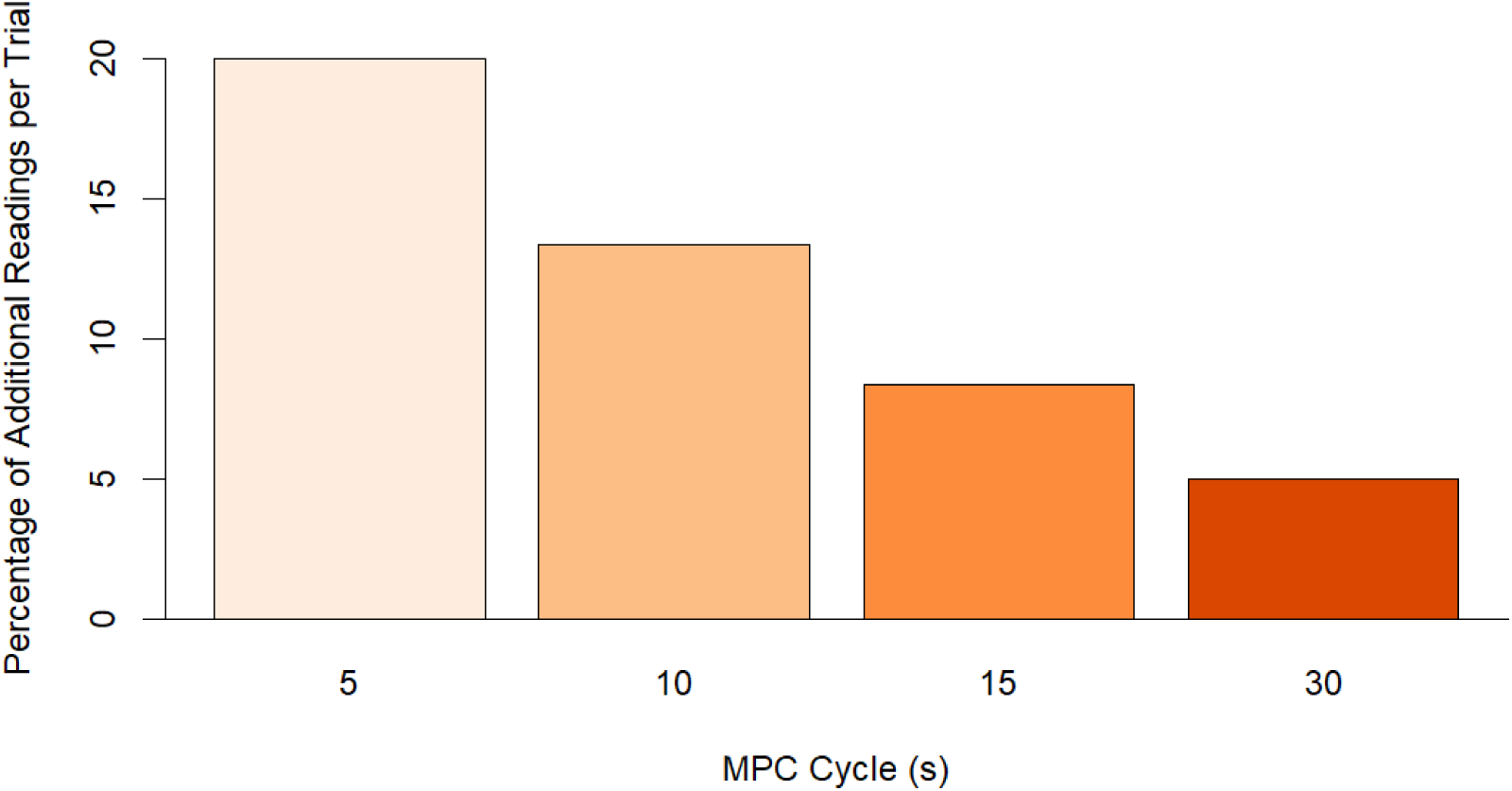
Percentage of additional readings per trial by MPC cycle length.

Supporting our first hypothesis that the number of additional readings will increase proportionally as MPC cycle length is reduced, no significant difference was found between the actual and expected proportions of additional readings for cycles of 5 seconds (χ^2^=0.56, df=1, p=0.46), 10 seconds (χ^2^=0, df=1, p=1), or 30 seconds (χ^2^=0, df=1, p=1). This shows that the actual proportion of additional readings for the 5, 10, and 30 second cycles was not statistically different from the expected proportions of 25%, 12.50%, and 4.17% respectively, calculated proportionally based on data from the 15 second cycle. Supporting our second hypothesis that an equal number of Type A and B errors will be present, no significant difference was found between the proportion of type A and B errors in trials of MPC cycle length 5, 10, 15, or 30 seconds (χ^2^=0, df=1, p=1; χ^2^=0, df=1, p=1; χ^2^=0, df=1, p=1; χ^2^=1.37, df=1, p=0.24). Supporting our third hypothesis that each ’unknown’ additional reading corresponds to specific antennas based on the type of error which has occurred, 100% of unknown additional readings (n=27) matched the expected antenna as outlined in Figure 5.

These results provide strong evidence backing up our proposed origin of additional readings generated by this system. Therefore, these ‘unknown’ or duplicate data points found in the csv file likely do not correspond to an actual transponder movement but are instead an artefact of the MPC cycle. Importantly, we can now interpret these additional readings and, thereby, increase data quality. For the following analysis of the trials of 60 data points, these additional readings were removed by hand. However, this method is not feasible regarding large scale data collection. Further, consistent discrepancies between the actual and recorded time of movement were observed, with the recorded time on average 120 seconds after the actual movement. Therefore, we created a Python script removing these additional readings and time discrepancy for raw data processing (https://github.com/BristolBeeGroup/BeeDrifting). This script also corrects for ‘unknown’ errors for which the correct reading can be predicted, improving data readability. The Following are rules used for this Python script:

To correct the time discrepancy:

▪ The timestamp shown should be lowered by 120 seconds for each data point To correct additional readings:

▪ To correct a type A error, ‘Unknown’ values one MPC cycle duration (± 0.5 seconds) after a directional movement from the same ID should be removed.

▪ To correct a type B error, ‘Unknown’ values one MPC cycle duration (± 0.5 seconds) before a directional movement from the same ID should be removed.

▪ To correct a type C error, ‘Arriving’ values one MPC cycle duration (± 0.5 seconds) after an ‘Arriving’ movement from the same ID should be removed.

▪ To correct a type C error, ‘Departing’ values one MPC cycle duration (± 0.5 seconds) after a ‘Departing’ movement from the same ID should be removed.

To correct ‘unknown’ errors for which the correct reading can be predicted:

▪ ‘Unknown’ readings belonging to an ID whose preceding reading is ‘departing’ at least 3 minutes prior, and whose proceeding reading is ‘departing’ at least 1 minute after, should be changed to ‘arriving’. This is based on the assumption that foraging trips last at least 3 minutes and in-hive stays last at least 1 minute.

▪ ‘Unknown’ readings belonging to an ID whose preceding reading is ‘arriving’ at least 1 minute prior, and whose proceeding reading is ‘arriving’ at least 3 minutes after, should be changed to ‘departing’.

To identify ‘drifting’ bees:

▪ Any ID that is recorded by multiple readers should be flagged as a ‘drifter’.

### (b) Accuracy of the system

#### Test 1 - MPC cycles

The percentage accuracy of the system decreased with decreasing MPC cycle length and was greatest when using a 30 second cycle (100%), followed by 15 (98.33%), 10 (96.67%), and 5 second cycles (93.33%) (Figure 19). However, no significant difference in percentage accuracy was observed between any of the 4 groups (χ^2^=5.15, df=3, p=0.16). Despite this non-significant result, the 15 second cycle was chosen as the default condition to use in subsequent tests due to its high accuracy but short duration promoting ease of experimentation.

**Figure 19:**
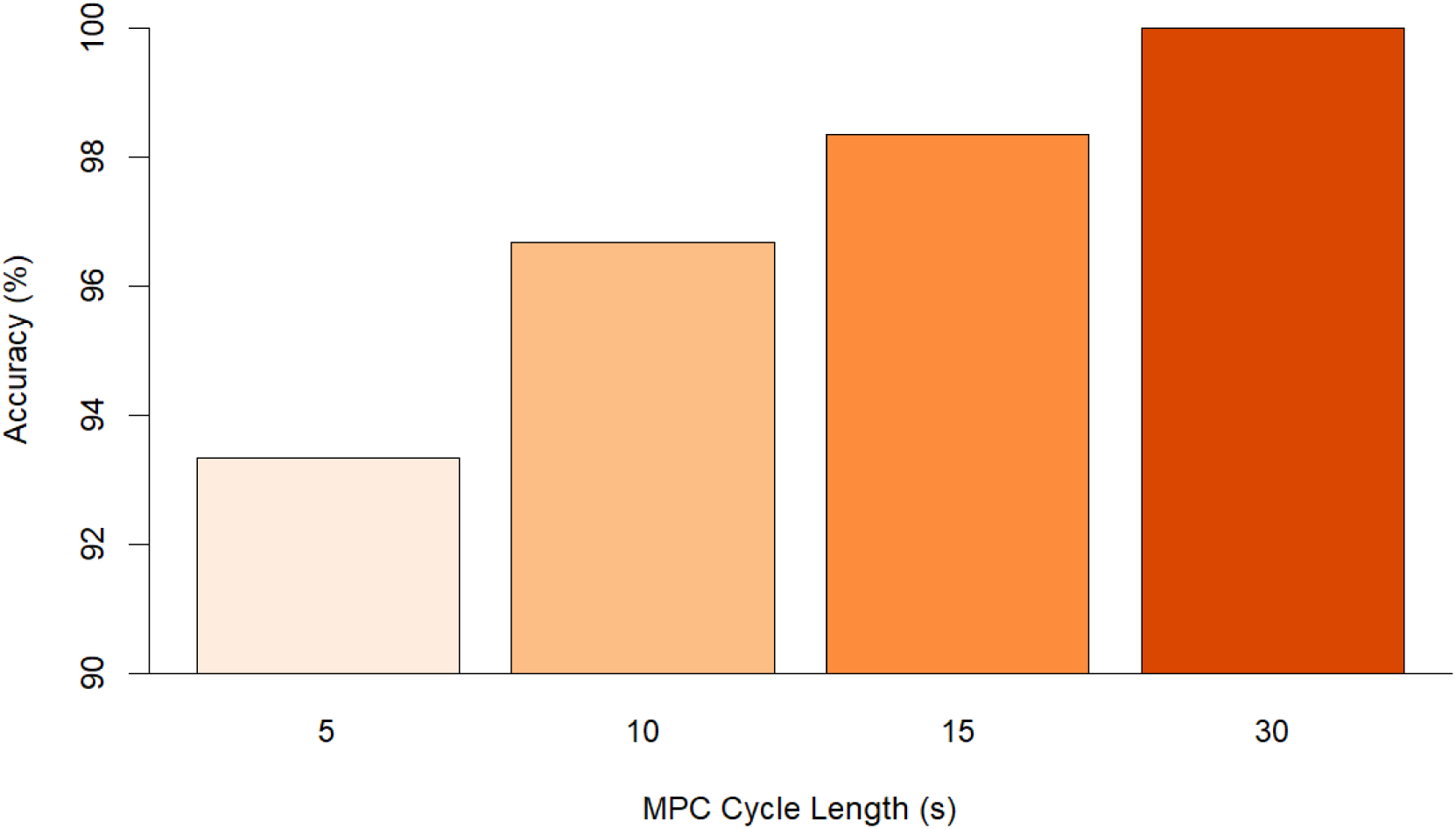
Accuracy rate of RFID System by MPC Cycle Length.

#### Test 2 – Dead bee

No significant difference was observed between the ‘dead bee’ strip and the foam bee strip (χ^2^<0.0001, df=1, p=1), suggesting that honeybee physiochemistry had no impact on transponder detection.

#### Test 3 - Speed of movement

No significant difference was observed between any of the 3 movement speeds (χ^2^=1.01, df=2, p=0.60), suggesting that the speed of the transponder passing through the reader has little to no impact on transponder detection, within a feasible range.

#### Test 4 – Transponder orientation

When using a ‘foam bee’ strip, the upside-down transponder trial produced a significantly lower accuracy than the upright transponder (χ^2^= 6.41, df=1, p<0.05; Figure 20A). However, no significant difference was observed when using the ‘dead bee’ strip (χ^2^<0.0001, df=1, p=1; Figure 20B). The reason for this discrepancy between the ‘foam bee’ and ‘dead bee’ strips is unknown, but the significant result for the ‘foam bee’ trial highlights the importance of correct application of transponders onto honeybees. They must be positioned upright to generate the highest accuracy.

**Figure 20:**
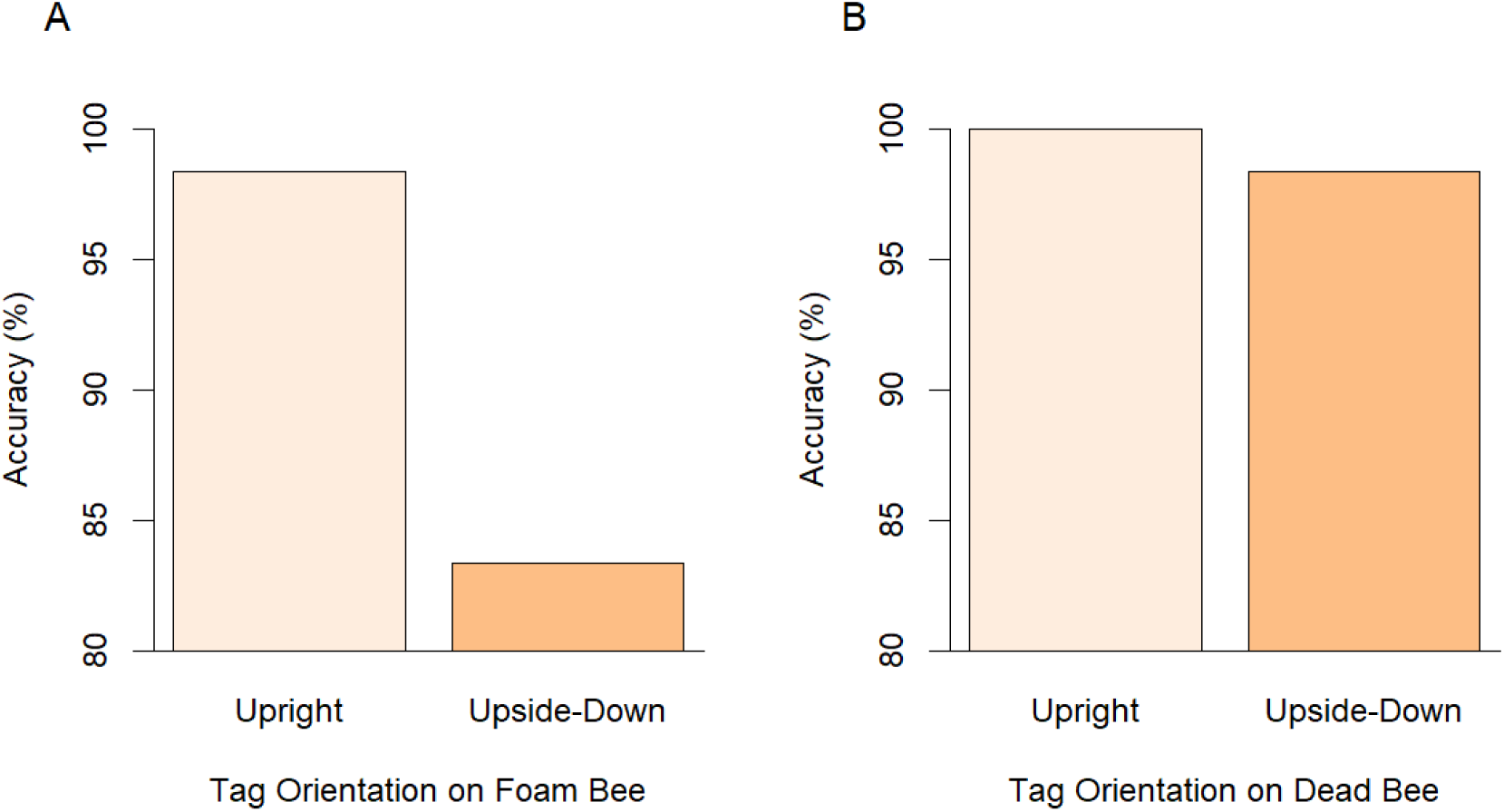
Accuracy rate of RFID System by transponder orientation using (A) the ‘foam bee’ strip; (B) the ‘dead bee’ strip.

#### Test 5 – Position in reader

When using the ‘foam bee’ strip, no significant difference was observed between the bottom and the top (χ^2^<0.0001, df=1, p=1; Figure 21A). However, both the left and right sides produced a significantly lower accuracy than the bottom (χ^2^=7.500, df=1, p<0.01; χ^2^=4.32, df=1, p<0.05). When using the ‘dead bee’ strip, no significant difference was observed between the bottom and the top or the right (χ^2^<0.0001, df=1, p=1; χ^2^<0.0001, df=1, p=1; Figure 21B). However, the left side was significantly less accurate than the bottom (χ^2^=12.42, df=1, p<0.0001). These results evidence that the left, and potentially right, sides of the reader produce a lower accuracy of readings compared to the other sides. Whilst this may instead be caused by the 90° orientation of the transponder rather than position in the reader, the outcome is the same, and experimenters should take care to determine whether this must be controlled for their experiment, dependent on the frequency of honeybees travelling on the side wall.

**Figure 21:**
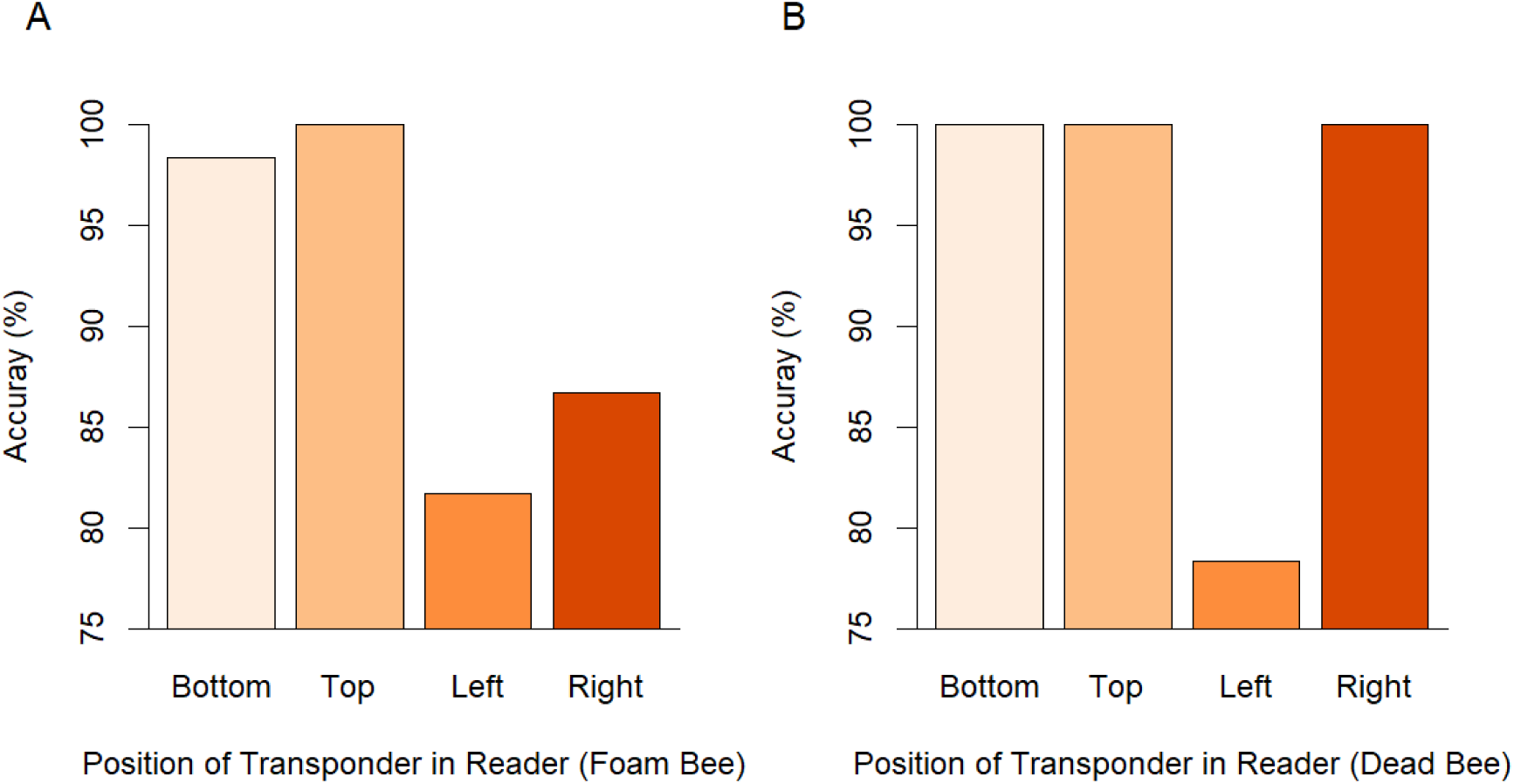
Accuracy rate of RFID System by position of the transponder in the reader for (A) the ‘foam bee’ strip; (B) the ‘dead bee’ strip.

#### Test 6 - Effectiveness of the second reader

No significant difference was observed between reader 1 and reader 2 for either ‘foam bee’ strips (χ^2^<0.0001, df=1, p=1), or ‘dead bee’ strips (χ^2^<0.0001, df=1, p=1), suggesting low variation between the 2 readers.

#### Test 7 - Distance from reader

No significant difference was observed between the 12mm reference and 9mm or 15mm (χ^2^<0.0001, df=1, p=1; χ^2^<0.0001, df=1, p=1; Figure 22). However, the accuracy of 18mm was significantly lower than the reference of 12mm (χ^2^=49.23, df=1, p<0.0001). This has little implications for the standard use of this apparatus with honeybees, as the average distance of a transponder from the reader will be ∼12mm. However, these results highlight the need for an enclosed entrance to the reader of maximum height 15mm to maximise accuracy.

**Figure 22:**
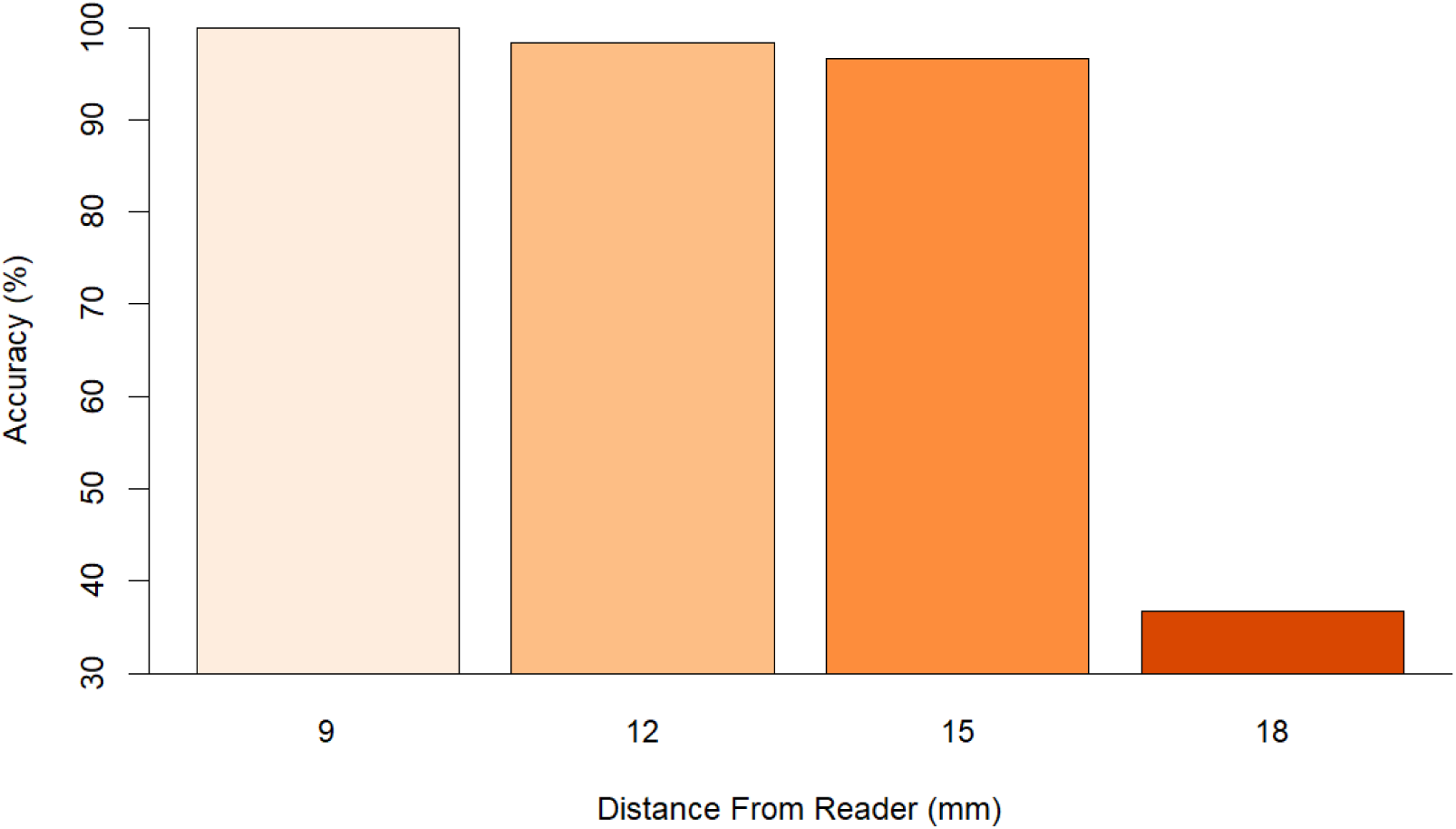
Accuracy rate of RFID System by distance from the reader.

#### Test 8 – Multiple transponders

A total of 300 data points were collected for the 5-transponder group, and 600 for the 10-transponder group. No significant difference was observed between a single transponder and 5 transponders through the reader simultaneously (χ^2^=0.26, df=1, p=0.63; Figure 23). However, the accuracy of 10 transponders was significantly lower than a single transponder (χ^2^=26.97, df=1, p<0.0001). This significant result likely has few implications for study’s employing this apparatus, as it is unlikely that greater than 5 tagged honeybees will pass through the reader at the same time. However, potential risk of accuracy loss can be mitigated by restricting the proportion of honeybees which are tagged.

**Figure 23:**
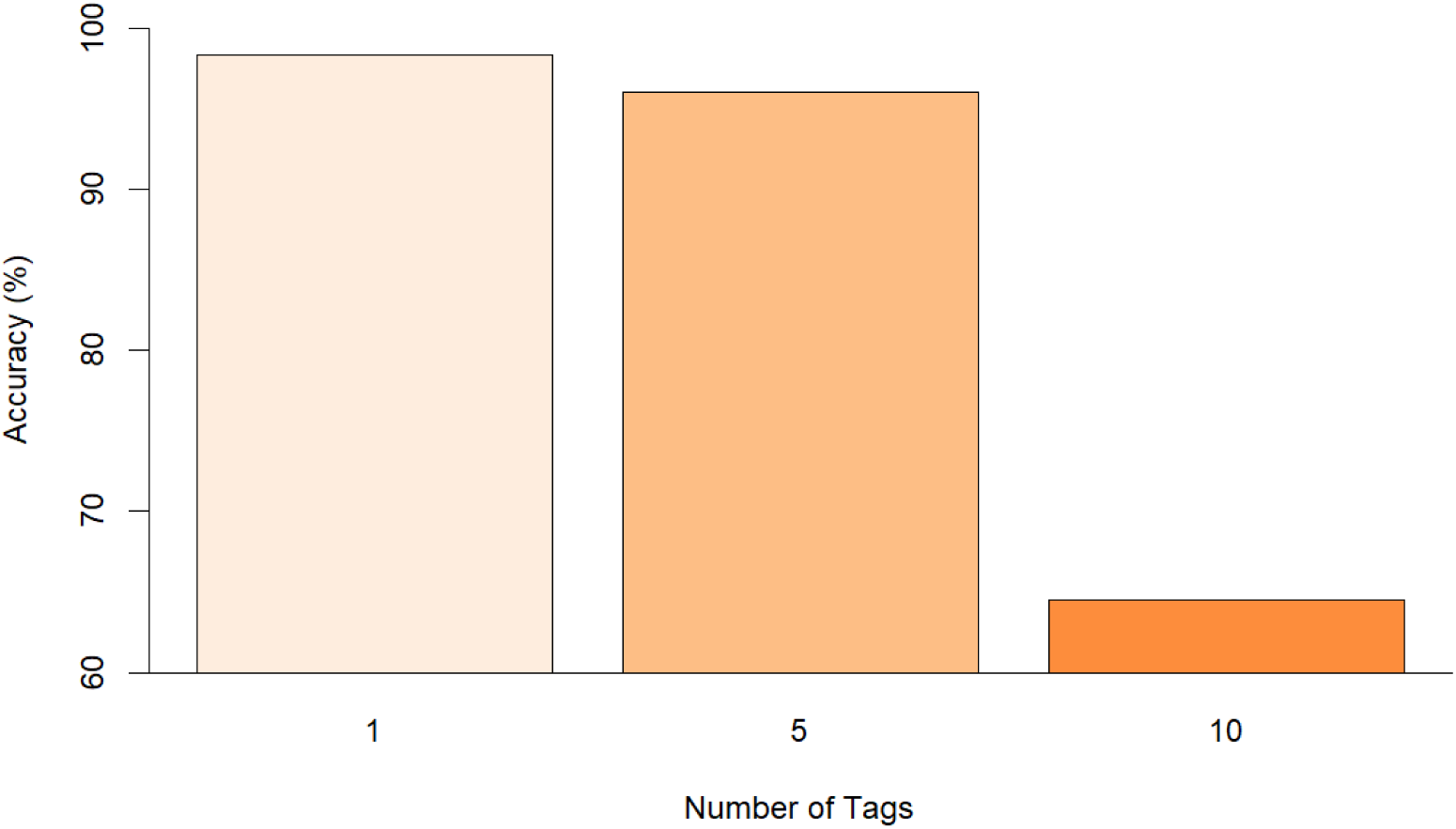
Accuracy rate of RFID System by number of simultaneous transponders.

#### Test 9 – Visual tag

No significant difference was observed between transponders with added number tags and either ‘foam bee’ strips (χ^2^=0.26, df=1, p=0.61), or ‘dead bee’ strips (χ^2^<0.0001, df=1, p=1), suggesting that visual tags can be used on top of RFID transponders without concern about decreased accuracy.

#### Test 10 – Readers close together

Whilst the accuracy of the readers 15cm apart was 3.3% lower than when the readers were 160cm apart, this was not significant (χ^2^= 0.26, df=1, p=0.61), suggesting that the distance between the readers has minimal effect. However, based on the manufacturer’s guidance, it is advisable to maintain a minimum of 150cm between individual readers.

#### Test 11 - Effects of water soaking

No significant difference was observed between transponders soaked for 5 minutes in water and those that weren’t (χ^2^<0.0001, df=1, p=1), suggesting that transponders are likely to operate effectively even when wet, eliminating concerns the impact of rain on system efficacy.

#### Test 12 – Effects of painting RFID transponder

No significant difference was observed between painted transponders and non-painted transponders (χ^2^<0.0001, df=1, p=1), evidencing painting the transponders as a viable method of individual identification.

### (c) Field testing

A total of 3 individual honeybees (of 18) were identifiable at the feeder as they possessed either painted RFID transponders (2) or an RFID transponder with a visual tag (1) from a previous experiment. Towards the start of the observation period, foraging trips were less frequent but lasted for longer, potentially owing to the need for ‘waggle dancing’ to recruit more honeybees to the feeder. As more honeybees were recruited to the feeder, the length of foraging trips decreased, and the frequency increased. Figure 24 shows all the observed foraging trips and visits to the colony recorded within the timeframe, collected by a human observer and the RFID system respectively.

**Figure 24:**
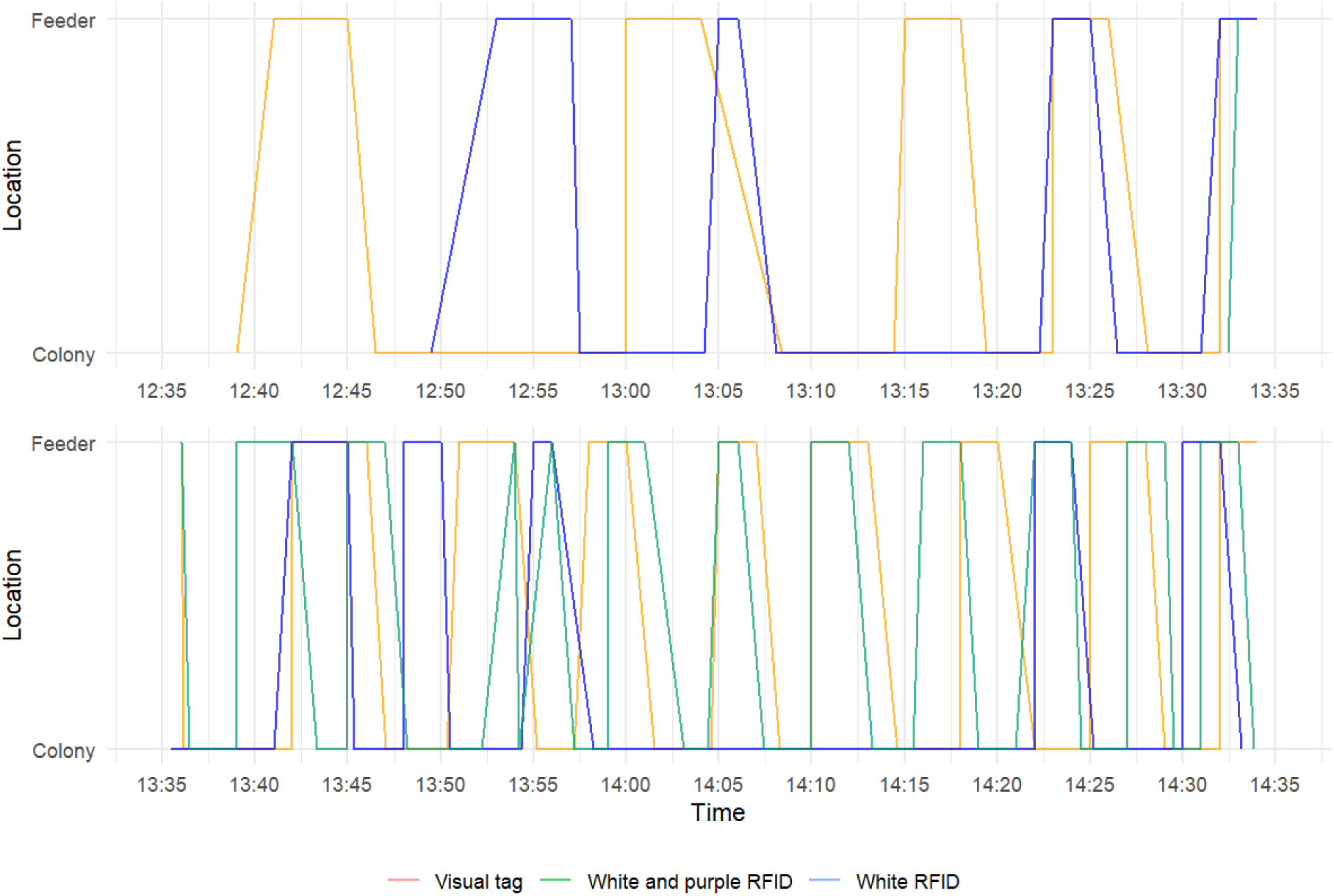
Foraging trips of 3 RFID-tagged honeybees over a 2 hour period between 12:35 and 14:35 pm: The feeder location reflects visual observations, and the colony location reflects RFID data. The individual with the visual-tag is represented by the green line; the white and purple RFID by the orange line; and the white RFID by the blue line.

For the honeybee tagged with a white and purple RFID transponder, a total of 13 observed foraging trips was recorded by the observer at the feeder. Correspondingly, the RFID system logged 13 departures and 13 arrivals at the colony, demonstrating full alignment with the direct observations. Similarly, a bee tagged with a white transponder completed 9 foraging trips, and an individual possessing an RFID transponder with a visual tag completed 11 foraging trips based on visual observations at the feeder, all of which were perfectly matched by RFID data. In all cases, the RFID system exhibited 100% efficiency in tracking the bees’ movements, with every departure and arrival detected. Based on this data, the system performed near perfectly, with RFID readings between foraging trips showing the honeybees returning to the colony, as would be expected. The only limitations of the RFID system presented as occasional ‘unknown’ errors rather than a directional movement. However, these can be interpreted correctly using our Python script (see above). No other technical difficulties arose from the use of the RFID system in combination with the observation hive.

However, the application of RFID transponders posed a challenge. Of the 18 RFID transponders attached to individual honeybees for this test (9 with number tags and 9 painted), only 3 (all painted) were later observed during the period of observation, 5 days after application of the numbered RFID transponders, and only 24 hours after application of the painted RFID transponders. Tagged foragers were observed attempting to remove their transponders immediately after tagging, and upon returning to their hive, other colony members were seen assisting them in tag removal. Potential solutions to this issue include tagging the bees shortly after hatching and re-introducing them to their home colony coated in sugar water as an appeasement, in addition to applying smoke to calm the

bees, wearing fresh gloves during tagging to reduce the impact of non-colony chemicals, and housing the tagged bees in a container partially joined to the hive to allow transfer of pheromones and acclimation. Such strategies are more difficult to impose on tagged foragers. Another possibility is to house the RFID transponders in a small container with a mesh lid, inside the colony prior to application, to allow transfer of colony-specific odours onto the transponder. Future efforts to tag honeybees with individually identifiable RFID transponders should use the strategies outlined above when introducing tagged bees to a colony. They should also use painted RFID transponders rather than RFID transponders with visual tags, as the former are more secure once attached, and offer the same service of identification.

Our successful use of this new RFID system for tracking honeybee foraging behaviour not only shows its promise when monitoring honeybee behavioural dynamics but also opens up new possibilities for researching a wide range of behaviours in other central-place foragers as small as ∼20mg due to the low mass (2.1mg) of the mic3®Q1.6 transponders. This will help to address the gap in the literature regarding the use of RFID technology to investigate behaviour in a wider range of *Hymenoptera*. By utilizing this RFID system to study the behaviour of smaller wasp and bee species, as well as larger ants, researchers can produce results which are not achievable using alternative methods such as direct observation, as RFID overcomes the issues of low sampling volume, visibility constraints, and limited observation times, allowing for a more comprehensive analysis of behavioural patterns in these species.

## 5. SUMMARY

Our study explored the efficacy of using an RFID tracking system for monitoring honeybee movements into and out of the hive. We confirmed that longer MPC cycles lead to a lower number of additional readings, though this was not significant, and a 15 second cycle length was suggested as optimal for our purposes. We investigated the cause behind additional readings, confirming that these do not represent actual transponder movements but are artifacts of how the system records transponder detections. Rules for a Python script have been proposed to automatically correct these errors for large-scale data collection, as well as correcting a time discrepancy, predictable ‘unknown’ errors, and identifying ‘drifters’. We investigated factors affecting the accuracy of the system, discovering that on the whole, the system has a high accuracy, yet the orientation of the transponder (upside-down) on the honeybee and its position in the reader (left and right walls) can significantly reduce accuracy. Additionally, the reader entrance tunnel must be restricted to 15mm height, otherwise a reduction in accuracy is observed. Other variables, such as honeybee physiochemistry, transponder speed, reader distance, environmental factors (e.g., water exposure), and the application of acrylic paint or a visual tag, had little or no impact on accuracy. Multiple transponders simultaneously passing through the reader only affected accuracy when the number exceeded 5, which is unlikely in most real-world scenarios. The RFID system performed without errors in field conditions, but the primary challenge was maintaining the attachment of RFID transponders to the honeybees, which often removed their transponders, or had them removed by colony members. This was particularly prominent when using RFID transponders with visual number tags. Painted RFID transponders showed better retention and are recommended for future studies in honeybees. These findings propose optimal RFID system parameters and ways of improving tag application methods to ensure reliable data collection in future honeybee behavioural studies. Further, we propose the use of this system to investigate the behaviour of central-place foragers as small as ∼20mg which were previously understudied using RFID.

## Acknowledgements

J.M. and C.G. were funded by the Biotechnology and Biological Sciences Research Council (BBSRC) (grant: BB/W001977/1). A.P.C. was supported by the SWBio Doctoral Training Partnership funded by the BBSRC (grant: BB/T008741/1).

